# Modulating Prefrontal Cortex Activity to Alleviate Stress-Induced Working Memory Deficits: A Transcranial Direct Current (tDCS) Study

**DOI:** 10.1101/2024.09.16.613190

**Authors:** Sumit Roy, Yan Fan, Mohsen Mosayebi-Samani, Maren Claus, Fatemeh Yavari, Thomas Kleinsorge, Michael A. Nitsche

**Affiliations:** Department of Psychology and Neurosciences, Leibniz Research Centre for Working Environment and Human Factors, Dortmund, Germany; International Graduate School of Neuroscience (IGSN), Ruhr University Bochum, Bochum, Germany; Department of Immunology, Leibniz Research Centre for Working Environment and Human Factors, Dortmund, Germany; German Centre for Mental Health (DZPG), Bochum; Bielefeld University, University Hospital OWL, Protestant Hospital of Bethel Foundation, University Clinic of Psychiatry and Psychotherapy

**Keywords:** tDCS, Stress, Working Memory, Brain Stimulation, EEG

## Abstract

This study explores the impact of stress on working memory (WM) performance, and the potential mitigating effects of transcranial direct current stimulation (tDCS) over the left dorsolateral prefrontal cortex (dlPFC) and ventromedial prefrontal cortex (vmPFC). The study had a crossover, randomized, single-blind, sham-controlled design, with stress induction as within-subject and stimulation condition as between-subject factors. We assessed stress-induced WM deficits using aversive video clips to induce stress and a verbal n-back task to assess WM performance. We analyzed physiological (cortisol and heart rate), behavioral, and electroencephalographic (EEG) changes due to stress before, during, and after WM task performance and their modulation by tDCS. Stress impaired WM performance in the sham stimulation condition for the 3-back load, but not for 2-back or 4-back loads in the WM task, and was associated with elevated physiological stress markers. tDCS over the vmPFC led to better WM task performance while stimulation over the dlPFC did not. Active tDCS with both dlPFC and vmPFC stimulation blunted cortisol release in stress conditions compared to sham. The EEG analysis revealed potential mechanisms explaining the behavioral effects of vmPFC stimulation. vmPFC stimulation led to a decreased P200 event-related potential (ERP) component compared to the sham stimulation condition and resulted in higher task-related alpha desynchronization, indicating reduced distractions and better focus during task performance. This study thus shows that the vmPFC might be a potential target for mitigating the effects of stress on WM performance, and contributes to the development of targeted interventions for stress-related cognitive impairments.

**Significance Statement:** This study investigates how stress impacts working memory performance and whether non-invasive brain stimulation can mitigate these effects. Using transcranial direct current stimulation (tDCS), we targeted specific brain areas in participants stressed by watching aversive videos. Stress reduced working memory performance and increased cortisol levels. However, tDCS applied to the ventromedial prefrontal cortex (vmPFC) improved task performance and blunted the cortisol response compared to a sham treatment. EEG data suggest that stimulation also enhanced focus during task performance and reduced emotional distraction. Our findings suggest that tDCS over the vmPFC could be a promising non-invasive brain stimulation method to counteract the negative effects of stress on cognitive functions.

## Introduction

Stress is a ubiquitous issue affecting the daily lives of millions of people worldwide (1). Identification of strategies to understand and combat stress is therefore relevant (2). Stress in classical terms is defined as a state of increased allostatic load resulting from exposure to various triggers, leading to a feeling of loss of control over physiological and psychological responses (3). Acute stress enhances the activity of the hypothalamus-pituitary axis (HPA), which shows a slow response, and the sympathetic nervous system (SNS) which shows a rapid response, causing increased levels of catecholamines, including norepinephrine, and cortisol, leading to the impairment of a broad range of cognitive processes, ranging from attentional control to WM performance (4, 5).

WM is defined as a system for temporarily storing incoming information and manipulating it according to task demands (6). WM processes are controlled by a large-scale network of cortical and sub-cortical areas which are prone to the effects of stress, leading to disruption of this system (7). It has been shown in numerous studies that WM performance is impaired by stress (8, 9).

Techniques such as EEG, which allows the exploration of brain oscillations and time-dependent changes in brain dynamics, can be used to investigate the physiological correlates of WM processes. Previous research has shown that theta oscillations control goal-relevant information and alpha oscillations suppress goal-irrelevant information during WM task performance (10, 11), while higher theta event-related synchronization (ERS) and higher alpha event-related desynchronization (ERD) have been linked to increased performance (12). Additionally, in the temporal domain, for ERPs, it has been shown that increased P300 activity is associated with efficient WM processing and memory updating (11, 13). The P300 amplitude is moreover sensitive to WM load and decreases with increased load (14). Furthermore, P200 activity has been linked to directing attentional resources to stimuli in WM tasks and decreases with WM task training (15, 16). Stress influences these EEG components by decreasing frontal theta (FT) activity (17), reducing the P300 component (18), increasing the P200 component (19, 20), and decreasing alpha ERD activity (21).

The dlPFC plays a crucial role in WM processes, and disruption of dlPFC activity leads to impaired WM task performance (22). Moreover, the dlPFC is highly susceptible to stress. Functional magnetic resonance imaging (fMRI) studies showed reduced activation of this area due to stress during WM task performance (23). Furthermore, the vmPFC is directly involved in stress, anxiety, and threat appraisal (24), and has relevant functional connections with the dlPFC and amygdala (25, 26). These connections are involved in emotional regulation and are impaired during stress (7). Stimulation of the dlPFC and vmPFC also influences emotional processing by reducing perceived valence and arousal during the presentation of emotional pictures (27), making these areas potential targets to ameliorate stress-induced WM deficits.

tDCS is a safe and non-invasive technique to modulate excitability, and activity of specific brain regions by delivering weak direct currents to the scalp. tDCS has been shown to alter cortical excitability during stimulation by modulating neuronal resting membrane potentials (acute effects) and induces neuroplastic effects via prolonged stimulation by glutamatergic synaptic mechanisms (28). tDCS also allows investigation of the causal contribution of brain areas and networks to psychological processes by exploring how stimulation modulates cognitive functions (29) and thus might be suited to improve understanding of brain and cognitive functions during stress. Anodal tDCS, which refers to a surface inward current over the target area, enhances cortical excitability, while cathodal tDCS, which refers to an outward current over the target area, results in excitability reduction with standard protocols at the macroscale level (30). Thus, the excitability-enhancing effects of anodal tDCS might be a potential tool to stimulate prefrontal areas to rescue stress-induced deficits in prefrontal cognitive processes.

Anodal tDCS over the left dlPFC has been shown to be effective in modulating working memory performance (31), and online tDCS (stimulation during task performance) seems to exert superior effects (32). Furthermore, anodal tDCS over right prefrontal areas has been shown to blunt cortisol response to stressors (33). Few studies have explored the impact of prefrontal tDCS on stress-induced WM deficits, with mixed findings (34–36). In the present experiment, focal tDCS montages to specifically stimulate the left dlPFC and vmPFC were used, and an aversive video stressor was applied. We collected and analyzed multiple data modalities to assess the effects of stimulation on physiological parameters and WM performance during stress, including cortisol, heart rate (HR), and EEG data.

Based on extensive work related to stress-induced WM deficits (8), we expected WM deficits caused by stress in the sham stimulation condition with stress also increasing physiological parameters such as cortisol, and HR (17, 23). Based on the roles of the vmPFC and dlPFC in emotional regulation and working memory performance, we anticipated that tDCS over these areas would improve WM task performance and limit stress-related cortisol release. Previous research has not extensively explored the associations of stress, WM performance, and tDCS together in the context of physiological markers like HR, thus the hypotheses that tDCS should ameliorate stress-induced HR increase are preliminary. Based on previous findings (17, 19), we furthermore expected a decrease of task-based FT activity and an increase of the P200 amplitude in the sham tDCS condition under stress, with stimulation reducing these alterations alongside improved performance. Given the significant roles of task-based parietal alpha and the P300 ERP component in WM task performance, we anticipated furthermore that tDCS would increase P300 activity and task-related alpha ERD in parietal areas.

## RESULTS

### Behavioral analysis of WM Task performance

#### Accuracy

Participants performed three blocks of the n-back letter task with different loads (n = 2, 3, and 4) in randomized order. Accuracy was calculated as the total number of correct responses (hits) made by participants for each n-back load. A 3-way repeated measures analysis of variance (rmANOVA) with emotional condition (control, stress), and WM loads (2, 3, and 4 back) as within-subject and stimulation groups (sham, dlPFC, and vmPFC) as between-subject factors were performed. The analysis revealed a significant main effect of load [F (2,144) = 360.1, p = <.001], with decreased accuracy with increasing load. Accuracy in the 2-back load condition was larger than in the 3-back and 4-back load condition, and the 3-back load condition resulted in higher accuracy than the 4-back load (all p<.001). A significant main effect of stimulation [F (2, 72) =3.9, p =.02] due to differences in task performance between stimulation conditions was also identified (please refer to table 1 for the results of this and the following ANOVAs). The vmPFC stimulation group outperformed the dlPFC (p = 0.009) and sham (p = 0.048) groups as shown by respective post hoc tests.

**Table 1.**
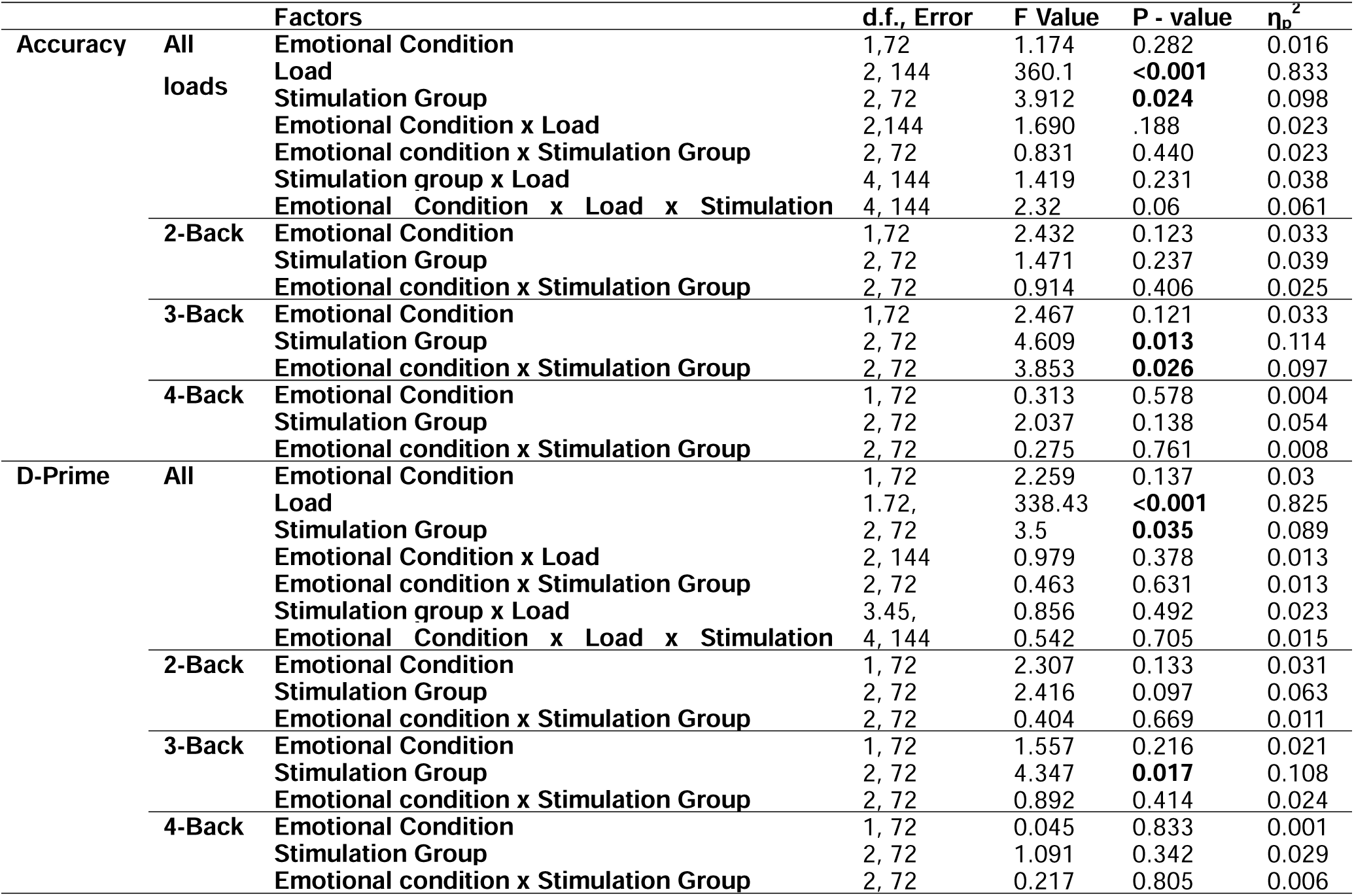
Main and interaction effects of the ANOVAs conducted for behavioral data of the WM task (accuracy and D-Prime). First, a 3-way RM ANOVA over all WM loads, and then secondary ANOVAs for each load were performed. Significant p-values are marked in bold. d.f. = degrees of freedom, η_p_2= partial eta squared.

Secondary 2-way rmANOVAs with emotional condition and stimulation groups as factors were conducted for each load separately. The 2-back and 4-back load ANOVAs revealed no significant main or interaction effects of emotional condition and stimulation. For the 3-back load condition, a significant main effect of stimulation [F (2, 72) = 4.6, p =.013] and a significant emotional x stimulation condition interaction [F (2, 72) = 3.85, p =.026] emerged. The post-hoc tests conducted for the 3-back condition revealed a decrease in the number of hits in the stress condition as compared to the control condition only during sham stimulation (p = 0.003), whereas no difference was found in the dlPFC and vmPFC stimulation groups between stress and control conditions. Post-hoc tests moreover revealed that vmPFC stimulation led to a larger number of hits as compared to dlPFC stimulation in the control (p = 0.03) condition, while the number of hits in the vmPFC group was larger than those in the dlPFC (p = 0.005) and sham groups (p = 0.002) in the stress condition (see Figure 1a).

**Figure 1.**
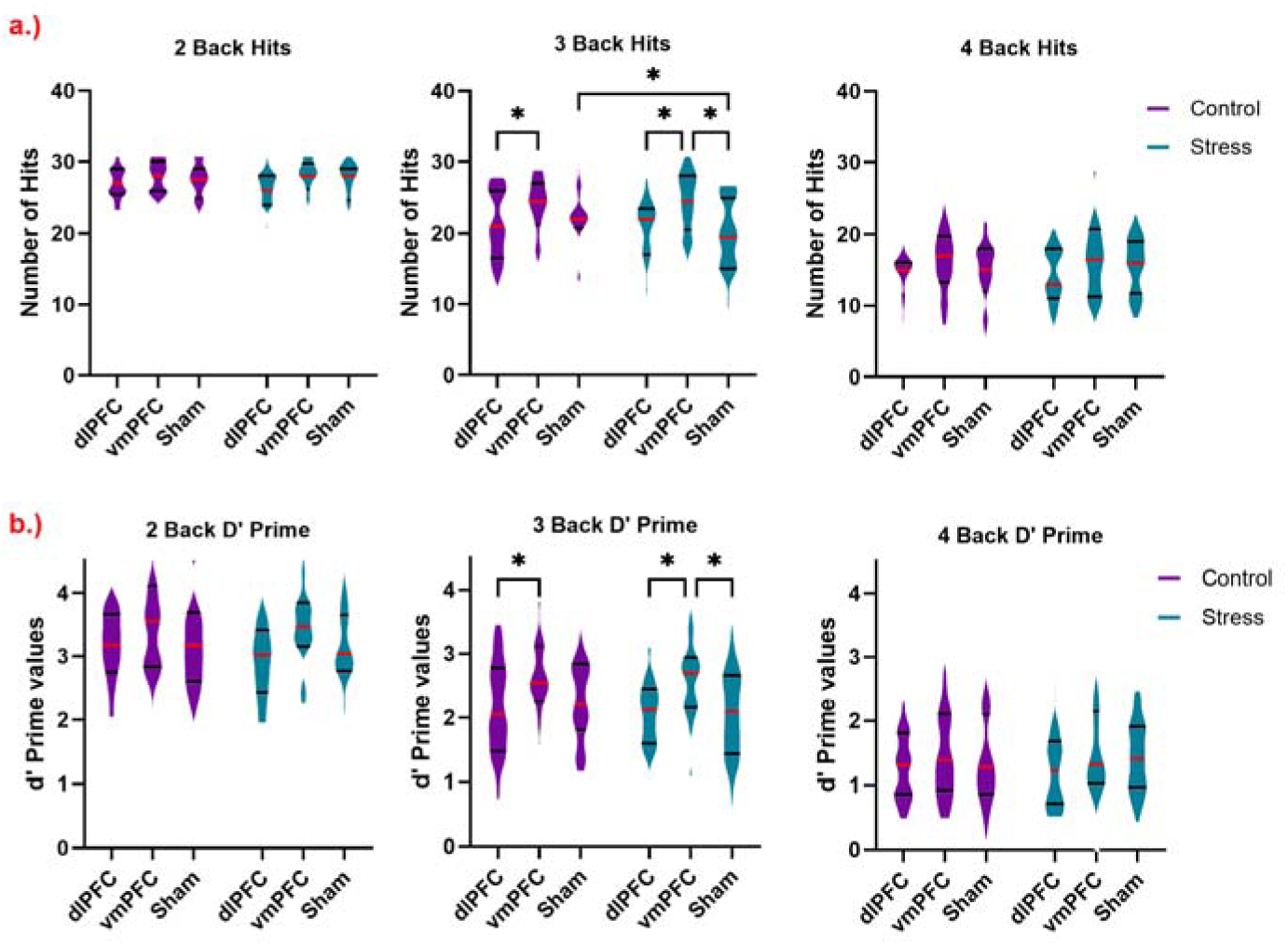
Behavioral data of WM task performance for accuracy and D-prime are shown. a.) Violin plots for the number of hits (accuracy) for all three n-back blocks are shown here. The highest possible number of hits in the task was 31. The y-axis shows the hit scores and the x-axis shows different tDCS groups (stimulation conditions) separated by emotional conditions (control, stress) colored violet and blue respectively. The median value is marked by horizontal red lines and quartiles by black lines for each violin plot. Significant differences are marked by asterisks (*) in black. b.) Violin plots for the D-Prime values for all three n-back blocks are shown here and plotted as described for accuracy.

#### D-Prime

D-prime as a performance index based on z-transform of hit rate and false alarm rate [z(hit rate) – z(false alarm rate)] (37) was calculated for each n-back load. As for performance accuracy, we performed first a 3-way ANOVA for all loads combined, and then secondary 2-way ANOVAs.

The 3-way rmANOVA showed significant main effects of WM load [F (1.7, 124.2) = 338.4, p <.001], and stimulation group [F (2, 72) = 3.5, p =.035]. Respective post hoc tests for WM load showed better scores in the 2-back compared to the 3-back and 4-back, and better scores in the 3-back compared to the 4-back load (all p <.001). Post-hoc tests conducted for the stimulation group showed stimulation-dependent differences in the D-prime level. tDCS over the vmPFC improved performance significantly as compared to stimulation over the dlPFC (p = 0.013), and trend-wise as compared to sham stimulation (p = 0.061) for both emotional conditions combined.

The secondary ANOVAs conducted for each n-back level separately revealed selective effects of stimulation and emotional condition on WM load. For the 2-back and 4-back WM conditions, no significant main or interaction effects of emotional condition and stimulation were revealed. For the 3-back task, however a significant main effect of stimulation emerged [F (2, 72) = 4.34, p =.017]. The post hoc tests revealed that vmPFC stimulation resulted in a larger score than dlPFC (p = 0.009) and sham (p = 0.017) stimulation for both emotional conditions. No significant main or interaction effect of emotional conditions was however identified (see Figure 1b.).

#### Physiological Measures of Stress

We analyzed cortisol, and HR to assess physiological changes due to stress and the influence of brain stimulation. Please refer to table 2 for a complete list of the ANOVA results for all physiological measures. For the results of an additional cytokine, and heart rate variability (HRV) data analysis, please refer to the supplementary table 6 and 7.

**Table 2.** Main and interaction effects of the ANOVAs conducted for physiological measures of stress (cortisol, and HR). Significant p-values are marked in bold, d.f. = degrees of freedom, η_p_2= partial eta squared.

| | Factors | d.f., Error | F Value | P - value | $\eta_p^2$ |
| --- | --- | --- | --- | --- | --- |
| Cortisol | Emotional Condition | 1,72 | 8.067 | **0.006** | 0.101 |
|  | Time | 2,144 | 12.32 | **<0.001** | 0.146 |
|  | Stimulation Group | 2,72 | 0.84 | 0.435 | 0.023 |
|  | Emotional Condition x Time | 1.74, 125.3 | 3.22 | **0.05** | 0.043 |
|  | Emotional condition x Stimulation Group | 2,72 | 2.7 | 0.07 | 0.07 |
|  | Stimulation group x Time | 4, 144 | 0.649 | 0.628 | 0.018 |
|  | Emotional Condition x Time x Stimulation Group | 3.48, 125.3 | 1.052 | 0.378 | 0.028 |
| HR | Emotional Condition | 1,74 | 0.578 | 0.449 | 0.008 |
|  | Time | 6.2, 465.6 | 33.48 | **<0.001** | 0.314 |
|  | Stimulation Group | 2, 74 | .459 | 0.634 | 0.012 |
|  | Emotional Condition x Time | 8.12, 600.8 | 1.36 | 0.207 | 0.018 |
|  | Emotional condition x Stimulation Group | 2,74 | 4 | **0.022** | 0.098 |
|  | Stimulation group x Time | 12.5, 465.6 | 1.021 | 0.429 | 0.027 |
|  | Emotional Condition x Time x Stimulation Group | 16.2, 600.8 | 0.5 | 0.95 | 0.013 |

#### HR

In the HR analysis, the rmANOVA showed a significant main effect of time [F (6.2, 465.6) = 33.49, p <.001] and a significant interaction of emotional conditions and stimulation groups [F (2, 74) = 4, p =.022]. Respective post-hoc tests revealed significant HR differences in the sham tDCS condition between emotional conditions (p = 0.009) where the stress condition resulted in elevated HR as compared to the control condition, but no such differences were found for the dlPFC (p = 0.25) and vmPFC (p = 0.82) stimulation groups (see Figure 2a.).

**Figure 2.**
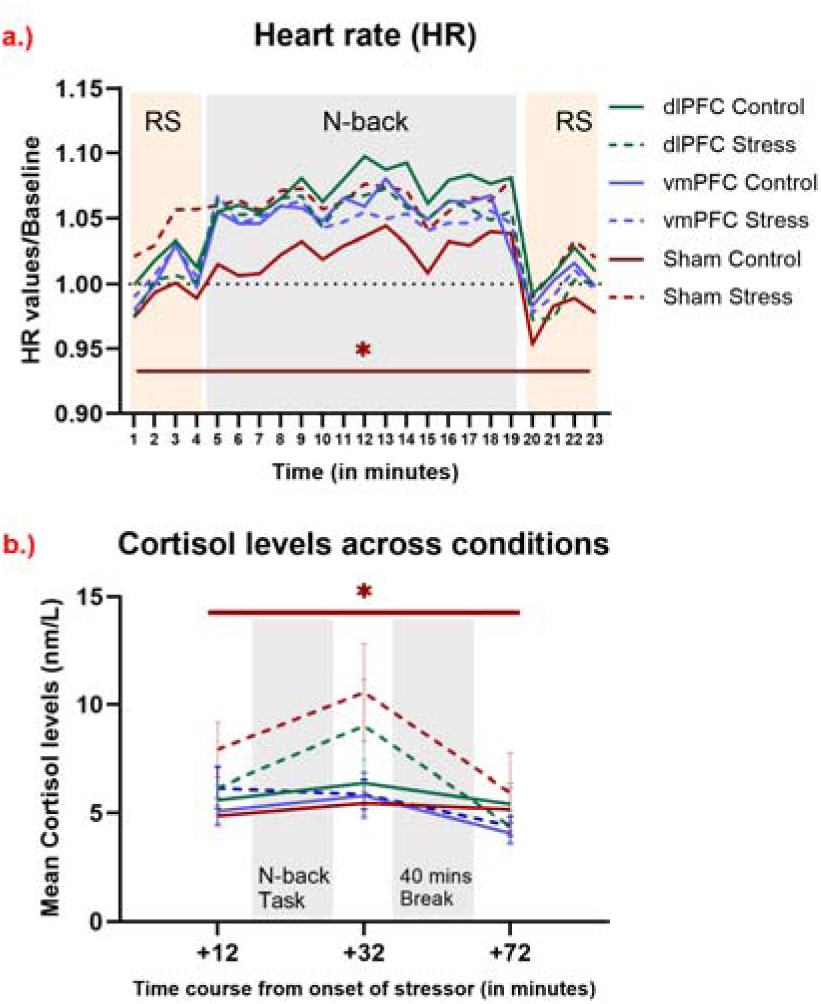
Physiological alterations induced by stress before, during, and after combined WM performance, and tDCS. a.) Heart rate before, during, and after task performance. Significant differences of HR between conditions (stress, control) were observed only in the sham condition, with increased HR in the stress as compared to the control condition for all time points, as shown by a red horizontal line marked by an asterisk (*). The y-axis denotes HR changes from baseline (first time point of the experiment as described in the methods section), and the x-axis denotes each minute of the experiment (from the end of the stress induction procedure). Resting-state (RS) and n-back task time frames are highlighted in the background with grey and orange colored watermarks (orange represents the resting state and grey the n-back task). b.) Cortisol levels before and after task performance are plotted. The y-axis displays mean cortisol levels (nm/l), and the x-axis shows the time course relative to the start of the movie clips. Error bars denote ± SEM. We found significant differences of cortisol concentration between conditions (stress, control) only in the sham condition for all time points, as shown by a red horizontal line marked by an asterisk (*).

#### Cortisol

The rmANOVA conducted for cortisol data revealed significant main effects of emotional condition [F (1, 72) = 8.06, p =.006] and time [F (2, 144) = 12.32, p <.001], as well as a significant emotional condition x time interaction [F (1.74, 125.3) = 3.22, p =.05]. As shown by the post hoc tests, in the stress condition cortisol levels were larger after task performance as compared to before (p = 0.036) and then decreased after the 40-minute break as compared to before task performance (p = 0.023). Furthermore, the interaction between emotional condition and stimulation group showed a trend-wise effect [F (2, 72) = 2.7, p =.07]. Respective post-hoc tests for all time points (before, after WM task performance, and after a 40 mins break) combined revealed a cortisol increase in the stress as compared to the control condition, which was significant in the sham tDCS condition (p < 0.001), while real stimulation reduced this response by diminishing cortisol level increases for both stimulation groups in the stress as compared to the control condition. No significant differences between control and stress conditions were observed for dlPFC (p = 0.392) and vmPFC stimulation for all time points, as done for the sham stimulation condition (p = 0.603) (see Figure 2b.).

#### EEG Results (3-Back)

We performed rmANOVAs for analyzing averaged P200 and P300 ERP components, task-based theta ERS, and task-based alpha ERD values (see Fig 3, Table 3). We also performed Montecarlo-based permutation statistics with cluster

**Figure 3.**
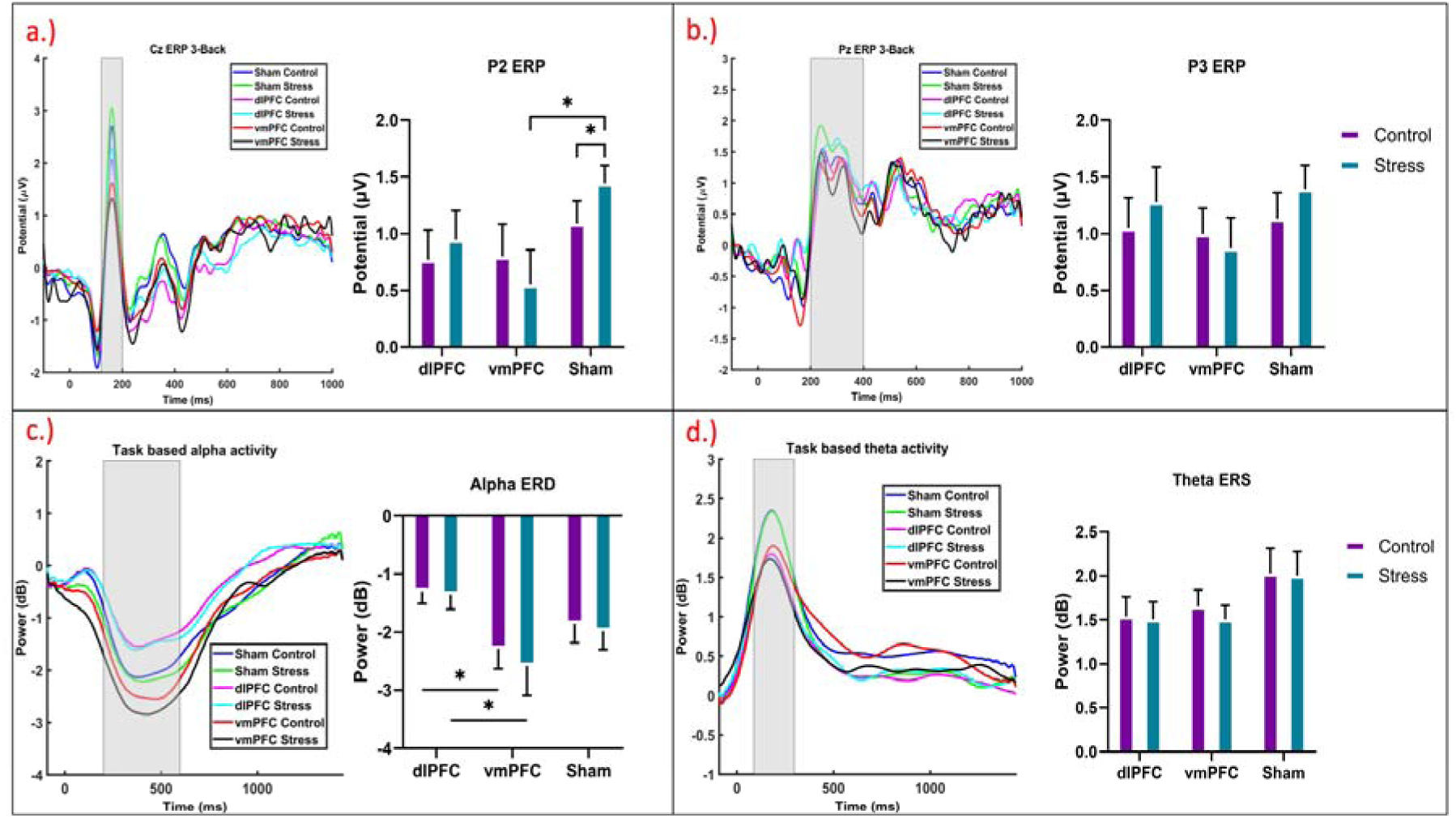
EEG results for the 3-Back load condition. a.) averaged ERP for all groups and conditions for EEG channel Cz. The x-axis represents time (in ms) and the y-axis shows the amplitude (µV). The time range of interest of the P200 (120-200ms) component is highlighted by a grey box. Respective averaged amplitude values were used for the statistical analysis, and shown as a bar plot next to it. The bar plot shows the averaged P200 component values for all groups and conditions with the x-axis indicating the groups and the y-axis the potential amplitude (µV). Significant differences are marked by an asterisk (*) in black color. b.) averaged ERP for all groups and conditions for the EEG channel Pz and the P300 (200-400ms) component amplitude, depicted as in a.). c.) averaged task-based alpha activity from parietal channels (Pz, P1, and P2) for all groups and conditions, the x-axis indicates time (ms), and the y-axis power (in dB). The time range of interest for ERD (200-600 ms) is marked by a grey box, and the mean power value within this time frame is plotted in a bar graph with the x-axis indicating groups, and y-axis power (dB). Significant differences are marked by an asterisk (*) in black color. d.) averaged task-based theta activity derived from frontal channels (Fz, F1, and F2) for ERS (80-300ms) plotted as for alpha ERD. Error bars denote ± SEM.

**Table 3.** Main and interaction effects of the ANOVAs conducted for EEG indices of the 3-Back load condition (P200 component, P300 component, theta ERS, and alpha ERD values). Significant p-values are marked in bold, d.f. = degrees of freedom, η_p_2= partial eta squared.

|  | **Factors** | **d.f., Error** | **F Value** | **P - value** | **$\eta_p^2$** |
| --- | --- | --- | --- | --- | --- |
| **P200 ERP** | **Emotional Condition** | 1,72 | .95 | .33 | .013 |
|  | **Stimulation Group** | 2,72 | 1.66 | .2 | .044 |
|  | **Emotional Condition x Stimulation Group** | 2,72 | 3.421 | **.038** | .087 |
| **P300 ERP** | **Emotional Condition** | 1,72 | 1.32 | .25 | .018 |
|  | **Stimulation Group** | 2,72 | .495 | .61 | .014 |
|  | **Emotional Condition x Stimulation Group** | 2,72 | 1.41 | .25 | .038 |
| **Theta ERS** | **Emotional Condition** | 1,72 | .57 | .453 | .008 |
|  | **Stimulation Group** | 2,72 | 1.6 | .2 | .044 |
|  | **Emotional Condition x Stimulation Group** | 2,72 | .202 | .818 | .006 |
| **Alpha ERD** | **Emotional Condition** | 1,72 | .99 | .323 | .014 |
|  | **Stimulation Group** | 2,72 | 3.14 | **.049** | .08 |
|  | **Emotional Condition x Stimulation Group** | 2,72 | .202 | .818 | .006 |

correction for task-based theta and alpha activity, (please refer to the supplementary material for the cluster results (Fig S6, Table S10)).

#### P200 ERP Component

The rmANOVA revealed a significant interaction effect of emotional condition and stimulation group [F (2, 72) = 3.42, p =.038] for the P200 ERP amplitude, but no significant main effects of the emotional condition and stimulation. The post-hoc analysis revealed an increased P200 amplitude in the stress condition compared to control for the sham stimulation group (p = 0.037), and the vmPFC group showed a reduced P200 amplitude compared to sham for the stress condition (p = 0.013).

#### P300 ERP Component

The respective rmANOVA did not reveal significant main or interaction effects of emotional condition and stimulation group on the P300 ERP component.

#### Theta ERS

In the rmANOVA, we found no significant main or interaction effect of emotional condition and stimulation group on task-based theta ERS activity.

#### Alpha ERD

The rmANOVA results revealed a significant main effect of stimulation group [F (2, 72) = 3.14, p =.049] on alpha ERD activity, but no significant main effect of emotional condition or a significant interaction effect. The post-hoc analysis revealed decreased power (i.e. higher ERD) in the vmPFC group as compared to the dlPFC group for both control (p = 0.022), and stress (p = 0.025) emotional conditions.

#### Correlations

We computed multiple simple correlations to understand the association of EEG and other physiological indices with Hits, and D-Prime scores for each stimulation group and emotional condition separately. We performed these correlations for all n-back loads separately (see Figure 4). Please refer to the supplementary material for 2-back (Fig S10) and 4-back (Fig S11) WM loads.

**Figure 4.**
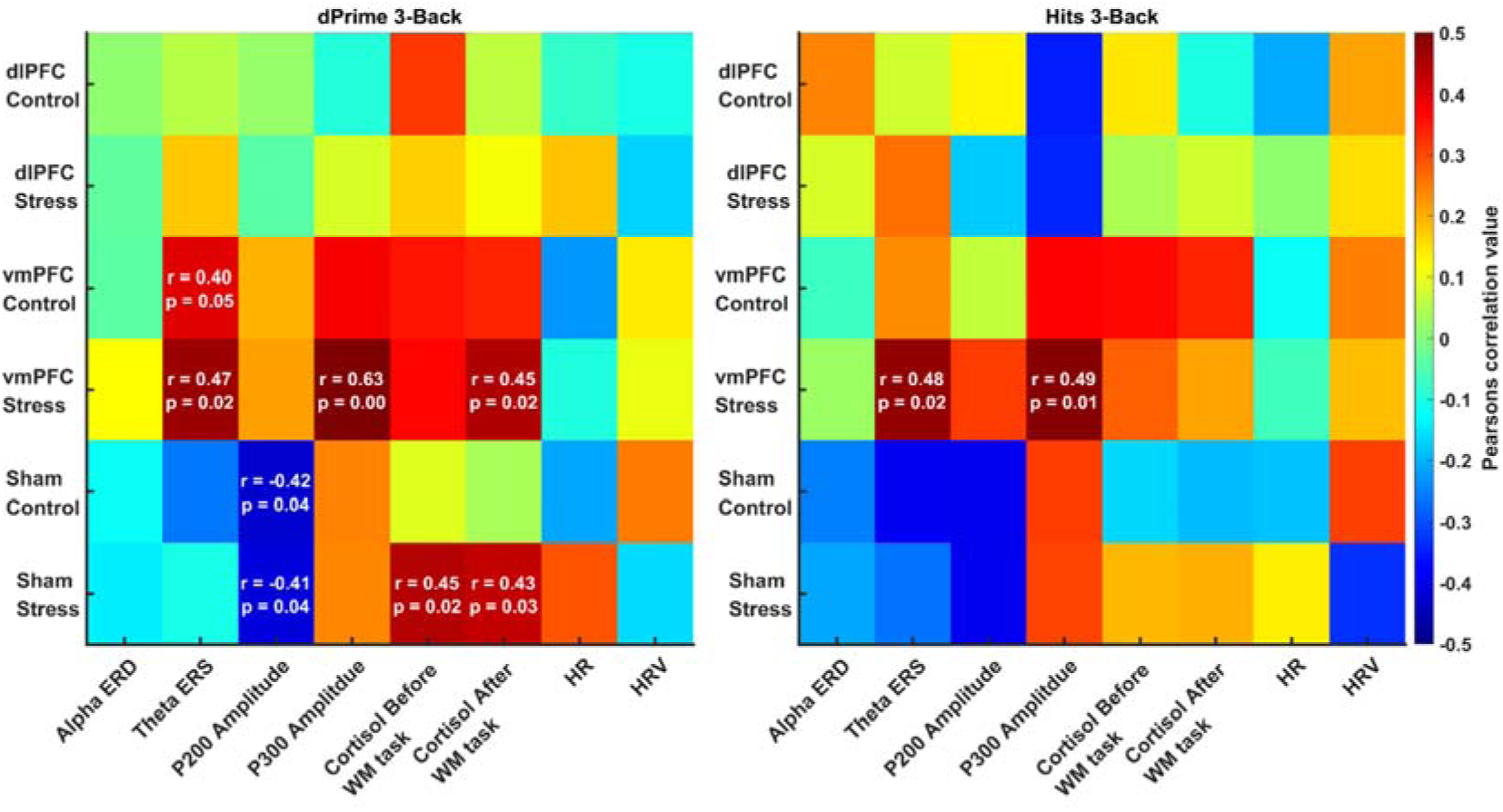
Heat map for Pearson’s correlation values in the 3-back load WM task for the EEG (alpha ERD, theta ERS, P200 amplitude, P300 amplitude) and other physiological indexes (cortisol levels before and after task performance, HR, and HRV during task performance) with the performance parameters Hits and D-prime for all stimulation and emotional conditions. The colors in the plot are showing r values with the color bar ranging from 0.5 to –0.5, red showing positive and blue showing negative correlations. For significant correlations, r-values, and p-values of the associations are shown.

#### 3-Back

In the 3-Back WM task, we found a significant association between the P200 amplitude and WM performance in the sham stimulation condition. Here, the P200 correlated negatively with D-prime under both emotional control and stress conditions. Moreover, theta ERS and the P300 amplitude were significantly positively correlated with both hits and D-prime scores in the vmPFC stress condition. Cortisol levels before and after task performance were positively correlated with D-prime scores in the sham stress condition and cortisol levels after task performance were positively correlated with D-prime scores in the vmPFC tDCS stress condition.

#### Side effects and Blinding

The results of the Chi-square tests indicated no significant heterogeneity of blinding in either the control (χ2 (2, N = 78) = 1.1, p = 0.57), nor stress condition (χ2 (2, N = 78) = .85, p = 0.655) with respect to stimulation condition.

For side effects, we found a significant effect of simulation group [F (2, 75) = 4.88, p =.01] on pain perceived during stimulation. The post-hoc tests showed that the vmPFC group reported less pain sensations as compared to the dlPFC (p = 0.037) and sham groups (p = 0.003). We also found a significant main effect of emotional condition [F (1, 2) = 4.18, p =.044] on nervousness, here the stress condition resulted in increased nervousness as compared to the control condition (p = 0.044) (see supplementary tables S2 and S3).

## Discussion

This study aimed to assess the effects of non-invasive brain stimulation (NIBS) on stress-induced WM deficits, including the assessment of the suitability and mechanistic role of stimulating different prefrontal regions in the context of stress. We used aversive video clips to induce stress as described in a previous study (38). In both, stress and an emotionally neutral control condition, participants had to perform an n-back task with different loads (2, 3, and 4 back) while receiving active or sham anodal tDCS over the vmPFC or dlPFC according to the respective intervention groups.

For the behavioral task analysis, we found a robust effect of stress on performance. Performance accuracy was reduced in the 3-back load condition in the sham tDCS condition, while no such effects were found for other WM loads. Interestingly, no significant effects of stress on D-prime levels in the sham tDCS condition were identified. Moreover, tDCS modulated WM performance, with stimulation over the vmPFC showing the largest effects. Stimulating over the vmPFC led to larger performance accuracy in the stress condition compared to sham and dlPFC stimulation, and also in the control condition, as compared to dlPFC stimulation for the 3-back load. A similar effect of stimulation was also seen for d-Prime performance, where the vmPFC tDCS group showed better performance in the stress condition compared to the sham and dlPFC stimulation groups in the 3-back WM task, and also in the control condition, as compared to dlPFC stimulation for the 3-back load.

These effects were only observed in the 3-back load condition, which corresponds to previous studies showing that stress impairs WM performance at high, but not low WM loads (39). Given that a 2-back load is relatively easy for young adults (40), we observed a potential ceiling effect in performance, whereas the 4-back load, being highly challenging, resulted likely in a floor effect. These findings suggest that these two load levels may not be sensitive enough for investigating the interaction between stress and stimulation. tDCS however rescued performance in the 3-back load condition. Since stress leads to downregulation of prefrontal activity (17, 23), most likely this effect can be mechanistically explained by a counteracting excitability-enhancing effect of tDCS. tDCS thus exerted promising effects on stress-related WM decline, and might be a valuable tool to achieve such effects. However, tasks that are too easy may not strain prefrontal processes to a sufficient level to be affected by tDCS, while tasks that are too difficult might not profit from tDCS due to floor effects.

A previous study has demonstrated beneficial effects of anodal tDCS over the right dlPFC on stress-induced WM deficits (35). Some other studies however reported no significant effect of anodal tDCS over the right dlPFC (34) and bi-frontal tDCS (36) on stress-induced WM deficits, thus leading to mixed results. It should be noted that all of these studies used conventional tDCS electrodes, thus resulting in stimulation of a broader prefrontal area (41), which might be not ideal for altering stress-or emotion-related effects on WM, since different prefrontal regions orchestrate emotions differently (42). In the present study, a more focalized approach was chosen to infer effects of specific prefrontal regions. Furthermore, the studies mentioned above used the Trier social stressor task (TSST) to induce stress. This might be problematic because of the arithmetic component of this stressor, which could affect a subsequent memory task by depleting cognitive resources, via deleterious effects of a prior task failure on performance in subsequent tasks (43).One study moreover utilized the arithmetic component of the TSST as working memory assessment to examine the role of tDCS, but they performed this task shortly after stress induction (34), while stress effects on prefrontal functions by HPA and SNS activity peaks at around 20 mins after stress induction (44). In our paradigm, we thus avoided introducing an arithmetic challenge before working memory task performance and assessed performance with a task not associated with that cognitive stressor and ∼20 mins after the start of the stressor.

The physiological findings support an impact of the stimulation on respective parameters. Cortisol and HR have been shown to be increased by stress (23). In our study, for cortisol levels, significant differences were revealed between stress and control conditions in the sham tDCS group showing stress-induced cortisol enhancement. This effect was however not observed in the active stimulation groups, indicating that stimulation controlled the release of cortisol and blunted this physiological stress response. Additionally, the sham group showed an increased HR in the stress compared to the control condition, a difference not seen in the stimulation groups. These results are in accordance with previous studies, which showed that prefrontal anodal stimulation blunted the release of cortisol after acute stress (33) and also reduced HR, however in a study not including acute stress induction (45). Prefrontal stimulation thus reduced stress-related upregulation of these physiological parameters. For stress-associated alterations of the cytokine profile, no substantial effect of stimulation or stress induction on cytokine levels was observed. In the present paradigm, only Interleukin-1 showing levels increased in the control as compared to stress condition (see Table S7).

We also collected online task-related EEG to shed light on the potential mechanisms underlying the observed improvements in WM performance during stimulation. Stimulating the vmPFC led to higher alpha ERD during task performance as compared to the dlPFC group for both emotional conditions and led to a decreased P200 amplitude during task performance in the stress condition as compared to sham stimulation for the 3-back load. In the sham condition, we observed increased P200 amplitudes and decreased FT activity in the stress condition from 910 ms to 1366 ms from stimulus onset (see figure S6) during 3-back WM task performance, aligned with previous findings that stress affects the early processing of stimuli impairing attention (19, 20) and FT activity affecting the early stages of WM updating (17). Behaviorally, this could have led to reduced accuracy (hit rate), while D-prime, which accounts for both hits and false alarms, remained unaffected. This suggests that reduced attention under stress not only lowered hits but also decreased false alarms, balancing D-prime despite the decline in accuracy. However, no such differences between stress and control conditions were seen in the real stimulation groups, indicating that stimulation mitigated these stress-related effects. Previous studies identified the roles of the P200 component and alpha ERD in cognitive processing under stress (19, 21). The P200 component, involved in early stimulus processing, is crucial for directing attentional resources to stimuli (19). The P200 has been shown to increase with stress and correlates with higher cortisol levels in males (46), indicating enhanced arousal. In our study, we also found that higher P200 amplitudes were negatively related to D-prime scores in both sham control and stress conditions for the 3-back task, linking higher P200 amplitudes to poorer performance in the sham group. Parietal alpha ERD suppresses irrelevant information and reduces distractions (47, 48) possibly caused by the aversive stressor in our study. vmPFC stimulation however led to higher alpha ERD. Our data suggest therefore that the lower P200 amplitude and larger alpha ERD most prominently observed for the vmPFC stimulation improved WM performance by reducing arousal and distractions, thus enhancing cognitive control and focus on the task.

While positive correlations of theta ERS and P300 activity with performance in the vmPFC stress condition were observed, P300 levels in the vmPFC stimulation condition were not significantly larger than in other groups. This is consistent with an involvement of the vmPFC not directly in WM processes (22) but rather in emotional responses, and the reduction of arousal resulting from emotional stimuli (24, 27). The foundation for this might be that the dlPFC is only indirectly connected with the amygdala, while the vmPFC has rather direct connections with this area, and acts as a mediator between the amygdala and the task-relevant dlPFC (26). The amygdala is known to influence emotional responses, and increased activation has been associated with decreased cognitive performance (49, 50). Specifically, the vmPFC diminishes emotional responses elicited by the amygdala (24) and loss of vmPFC activity leads to increased activation of the amygdala as response to an aversive stimuli (51).

Our results revealed a positive correlation between WM task performance, and cortisol levels after WM task performance in the vmPFC stress condition across all N-back loads (see figure 4, S10, S11). Moreover, in the Sham Stress condition, cortisol levels before and after the WM task performance correlated with each other in the 3-back load condition. This is consistent with previous research showing that increased cortisol responses to acute stressors can enhance WM performance in men (20). Interestingly, in our study, vmPFC stimulation led to a decrease in cortisol levels but still cortisol level correlated positively with performance. This suggests that vmPFC stimulation may have a dual effect: enhancing performance by strengthening WM-related networks while simultaneously reducing cortisol levels through better stressor recovery and control (24, 52)

Some limitations of this study should be taken into consideration. The sample population includes only males to control for effects of hormonal cycle on stress induction, thus limiting extrapolation of the results to other sexes. The stress induction paradigm was effective in eliciting stress, but might also include factors beyond pure stress, such as disgust due to the aversive content of clips. While the n-back task is a well introduced WM task, other WM tasks should also be explored to improve generalization of the findings. Future studies could also explore the impact of stress, including its modification by stimulation, on other cognitive processes like attention or decision making. Future studies using whole brain fMRI to identify the influence of stress on activation and connectivity of different cortical and sub-cortical areas as well as the impact of stimulation over prefrontal areas on respective fMRI parameters would be valuable to further clarify mechanisms. Specifically fMRI could provide information about the activation of medial brain areas like the amygdala and substantiate the influence of vmPFC stimulation on amygdala activity in the context of acute stress.

In conclusion, this study suggests a potential of NIBS for mitigating stress-induced WM performance deficits. We show that stimulation over the vmPFC led to better performance in a verbal n-back task as compared to sham and left dlPFC stimulation. Prefrontal stimulation also led to a blunted stress-related cortisol response suggesting physiological changes induced by stimulation. EEG analyses showed a reduced P200 amplitude and increased alpha ERD during task performance for the vmPFC stimulation condition, suggesting downregulation of the emotional response to the stressor as a probable cause for the impact of stimulation on performance.

## Methods

### Participants

Seventy-eight, healthy, right-handed male non-smokers, with handedness assessed by the Edinburgh handedness inventory (53), aged between 18-40 years (M = 25, SD = ±4.16) were included in the study. A medical check before the experiment was done to exclude participants with any chronic disease, psychiatric or neurological disorders, intake of any central nervous system (CNS)-acting medication or steroids, presence of any implants or devices in the body, oral inflammation, history of epilepsy, traumatic brain injury, and night shift working. A pre-experimental phone interview was done to make sure that the participants did not have any major emotional or physical trauma and no habit of watching extremely violent videos. Participants were divided randomly into three groups based on a generated random order (https://damienmasson.com/tools/latin_square/) when they arrived – anodal tDCS over the dlPFC, vmPFC, or sham stimulation. This study was approved by the Ethics Committee of the Leibniz Research Centre for Working Environment and Human Factors at TU Dortmund (IfADo) and aligns with the Declaration of Helsinki. Participants were informed about their right to withdraw from the experiment at any time and informed consent was taken before the start of the experiment. Participants were identical to a previously published study dedicated to the efficacy of the applied aversive stress induction protocol (38). The present study involves the WM part of that project. No group differences based on age, chronotype, or trait anxiety were found (Table S1).

### Stress Induction Procedure

Stress was induced using aversive video clips from the movie “Irreversible” by Gaspar Noe, while clips in the control condition were extracted from the movie “Comment j’ai tué mon père” by A. Fontaine. For a more detailed description of the stress induction methodology please refer to our previous study, which explored the effectiveness of stress induction via this protocol (38).

### Working memory task

We used three levels of the n-back letter task (n = 2, 3, and 4) in randomized order. Participants were seated in front of a computer monitor with a 50 cm eye distance. A pseudo-random set of 10 letters from (A-J) was presented. Each letter was displayed for 300 ms with a 1.7-second interval (blank screen). A different letter was displayed every 2 seconds. Letters were presented in black on a white background. Participants were required to respond with a keypress if the presented letter was the same as the letter presented 2, 3, or 4 (depending on the subtype of the task) stimuli previously. For each n-back level, 143 letters were presented, and a maximum of 31 correct responses (hits) could be achieved. The duration of a block was 5 minutes. The total duration of the task was approximately 16 min with 30 seconds rest between blocks. Participants were allowed to practice a short version of the task (with 46 letters) 3 times during the introductory session to reduce task learning effects and task-related stress. For behavioural outcomes from the task, we analyzed D-prime, accuracy, and reaction time (RT) for correct responses (for RT data, see Table S7).

### Stimulation Parameters

We employed computational modeling to determine the electrode montages for targeting the left dlPFC ansd vmPFC. A search space including all possible combinations of two square (5×5 cm) electrodes at 10-20 EEG positions, and various 4×1 electrode montages were explored via ROAST (https://www.parralab.org/roast/) (54). For each montage, the average electric field (EF) over the left dlPFC, right dlPFC, vmPFC, anterior cingulate cortex (ACC), and cerebellum was computed. The final montages for each of the two targets were selected by comparing the induced EF values achieved with different electrode configurations, to identify the electrode montages with the maximum average EF value in the target region, and minimum values in the other regions. This ended up in a montage with five round electrodes with a 1 cm radius for each of the target areas, as shown in Fig. 5. For vmPFC stimulation, the center electrode was positioned over the nasion, and four return electrodes were placed over F7, F8, Ex19, and Ex20. For targeting the left dlPFC, the center electrode was positioned over F3 and the four return ones were placed over Fp1, Fz, C3, and F7. Participants received stimulation during task performance (online) for approximately 17 minutes with 30 seconds of ramp-up and –down. Stimulation was applied with carbon rubber electrodes attached to the skull via Ten20 paste (Weaver and Co., USA). We applied Emla cream (AstraZeneca, UK) before stimulation at the electrode sites to reduce skin perceptions of stimulation. For sham stimulation, we applied a 30-second ramp-up, 30 seconds of stimulation, and 30 seconds ramp-down of stimulation intensity at the start and at the end to blind participants, no current was given during actual task performance. For all stimulation groups, the current intensity was 2mA.

**Figure 5.**
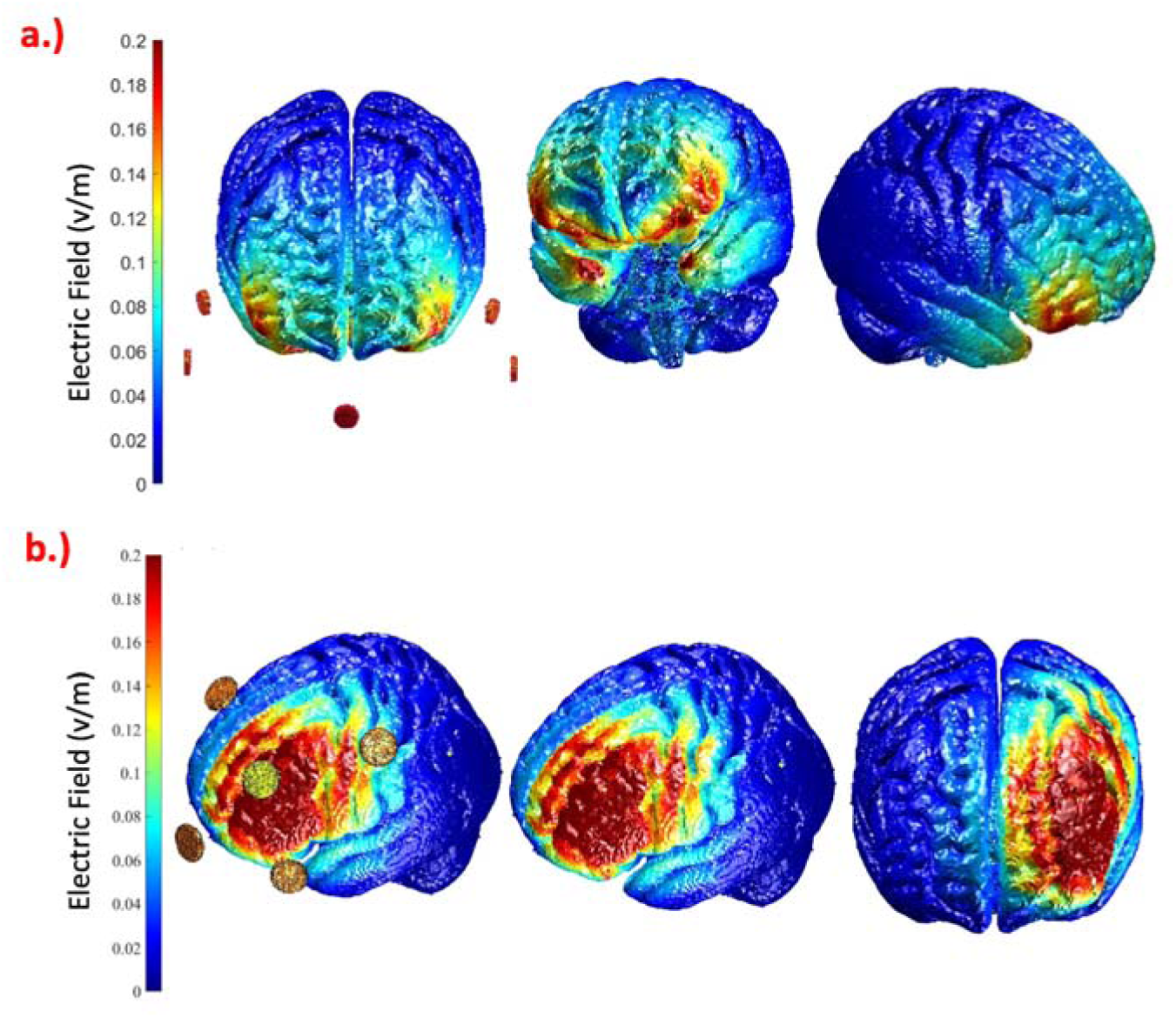
Simulated distribution of the electric field in the brain based on computational modeling for the selected electrode montage to target the (a) vmPFC with the electrodes over Nz (Nasion), F7, F8, Ex19, and Ex20, and (b) left DLPFC, with the electrodes over F3, Fp1, Fz, C3, and F7. The current amplitude was set to 2 mA, and all electrodes were discs with a 1 cm radius.

### EEG data Acquisition and processing

EEG was recorded during resting states (RS) and task performance against a reference electrode placed over the left mastoid, using sintered Ag-AgCl electrodes at 64 standard positions of the International on 10-20 EEG System (NeurOne, Finland). Electrode impedance was monitored and kept below 10 kOhm throughout the experiment. The sampling rate was 2000 Hz at an analog-to-digital precision of 24 bits. Offline EEG analysis was done using EEGLAB (v2022.1) (55), Brain Vision Analyzer (Version 2.2.0, Brain Products GmbH, Gilching, Germany), Fieldtrip (56), and custom MATLAB commands. Please refer to the supplementary material for the offline EEG data processing pipeline.

### ERP Analysis

ERP analysis was performed via the EEGLAB study function and custom MATLAB scripts. For ERP Data analysis, based on previous works on n-back task performance and stress, we analyzed the P200 ERP component (120-200 ms from stimulus onset) from the Cz channel (19, 20) and the P300 ERP component (200-400 ms from stimulus onset) from the Pz channel (11) for each task epoch which was then averaged over the epochs for each subject.

### Time Frequency (TF) Analysis

TF analysis was performed using the Morlet wave decomposition method implemented in EEGLAB STUDY using the newtimef function. 200 log-spaced frequencies in the range from 3 to 80 Hz, and 200 linearly spaced time bins for each task epoch were used for power calculation. Wavelet decomposition was done by using the parameters [3 0.8], where 3 cycles were used for the lowest frequency (3 Hz) and 16 cycles were used for the highest frequency (80 Hz). Power was normalized using decibel (dB) transform – (dB powerL=L10*log10 [power/baseline]). Based on previous works on WM performance and stress we analyzed Theta (4-8 Hz) ERS during the task from frontal electrodes (Fz, F1, and F2) for the time range 80 to 300 ms relative to stimulus onset (17), and Alpha (8-13 Hz) ERD activity during the task from parietal electrodes (Pz, P1, and P2) for time range 200 to 600 ms relative to stimulus onset (21). ERD and ERS of all epochs were then averaged over channels and epochs for each subject.

### Subjective Data

Two subjective self-report questionnaires were administered at various intervals during the experiment to obtain information about the subjective affective state of the participants: the positive and negative affect schedule (PANAS) (57, 58) and the State-Trait anxiety inventory-State (STAI-S). The PANAS obtains information about positive and negative emotions by 20 items, each rated on a 5-point scale from 1 (not at all) to 5 (very much). The STAI-S assessed state anxiety levels with 20 questions, rated on a scale from 1 (not at all) to 4 (very much). To assess the trait anxiety of the participants, we administered the State-Trait anxiety inventory-Trait (STAI-T) in the introductory session (see Experimental Procedure), which measures trait anxiety via 20 questions, rated on a scale from 1 (not at all) to 4 (very much) (59, 60). The morningness-eveningness questionnaire (MEQ) was also applied in the introductory session (61, 62). Participants were allowed to complete the questionnaires in either German or English, based on their preference. Data were collected via the online Sosci survey platform (63) and analyzed using custom MATLAB scripts.

### Side Effect and Blinding Questionnaire

After finishing each session, participants were asked to guess the stimulation condition (real/sham) and to rate side effects during (itching, tingling, burning, and pain) and after stimulation (skin redness, headache, fatigue, concentration difficulties, and nervousness) on a scale of 0 to 5, where 0 represents no and 5 represents extreme sensations.

### Experimental Procedure

Participants underwent a three-session experimental protocol, including a practice session aimed at reducing unspecific experiment– and context-related stress, conduction of a medical check, and training of the participants in WM task performance. Subsequently, two experimental sessions—control and stress— were administered, with session order randomized and counterbalanced according to a generated random order (https://damienmasson.com/tools/latin_square/). All experiments were conducted between 12 am and 6 pm to control for endogenous cortisol activity.

For the experimental sessions, participants arrived one hour before the core procedures of the sessions for EEG cap preparation, and tDCS and ECG electrode placement. The experiment began with the emotional intervention part described in a previous paper (38). After the emotional intervention, participants performed the n-back WM task for around 16 mins (3 blocks). During task performance, tDCS was applied. After task performance, salivary samples were taken, and PANAS and STAI were administered (see supplementary table 4), followed by an EEG RS recording (2 minutes with eyes open and 2 minutes with eyes closed). The experiment ended with a last saliva sample collection after 40 minutes and side effects and blinding questionnaire. This experiment comprised two parts: the first part, involving stress induction via movie clip presentation and assessment of the efficacy of stress induction, has been described in detail in a previous study of our group (38). The second part, which is described here, assessed WM performance with an n-back task during tDCS after movie clip presentation. Only the second part is presented here. Please refer to Figure 6 for a detailed overview of the procedure. Please refer to supplementary methods for HR, cortisol, and cytokine data acquisition and processing.

**Figure 6.**
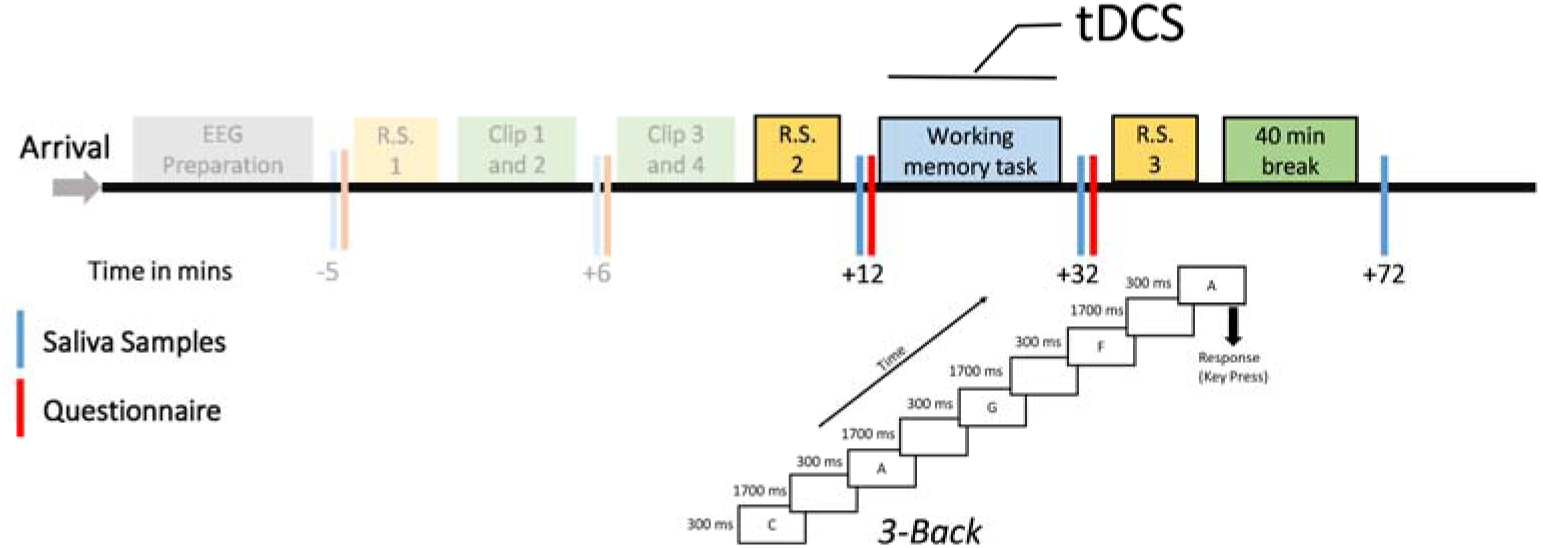
Experimental procedures: After the arrival of the participants, an EEG cap was prepared and the stress induction part was done as described in detail in a previous work (38). After resting state EEG (RS1), the movie clip presentation and another resting state EEG (RS2), participants performed the n-back task with three blocks (2, 3, and 4 back) in randomized order. The 3-back task is shown as an example, where the correct response (hit) is highlighted by a respective arrow. tDCS using small electrodes with a 1 cm radius was given in accordance with the stimulation group (sham, anodal dlPFC, and anodal vmPFC tDCS). After task performance, another RS EEG with 2 min Eyes Open (EO) and 2 min Eyes Closed (EC) conditions was performed (RS3). Blue bars show the time points of saliva (cortisol) sampling, and red bars show time points STAI and PANAS were administered. The time points for saliva sampling and questionnaire conduction in relation to the start of the intervention (clip presentation) are shown below the respective bars in minutes. HR (using bipolar electrodes positioned over the chest) and EEG were recorded during the whole experiment. The procedure was identical for control and stress sessions except for the kind of movie clips shown. The former part of the experiment, including the stress induction part, is shown in opaque color and was reported in detail elsewhere as it is beyond the scope of this article (38).

### Statistics

For data analysis, SPSS version 29.0.0 (IBM Corp., Armonk, New York, USA) was used, and data were plotted with GraphPad Prism 10. rmANOVAs for cortisol were performed with emotional condition (control, stress) and time (before, immediately after task performance, and after a 40 min break) as within-subject factors, and stimulation group as between-subject factor. Similar rmANOVAs were also conducted for HR with emotional condition and time (t1-t23) as within-subject, and stimulation group as between-subject factors. The first time point of the experiment before stress induction, as described in previous work (38), was taken as baseline, and then the change from baseline after emotional/control clip presentation to the end of the experiment was analyzed for HR data. For WM performance, based on outlier analyses, the WM data of three participants, for which individual means differed more than 2 standard deviations from the group mean were excluded from the analysis (one participant from the dlPFC, and two from the vmPFC group). rmANOVAs were conducted for accuracy (number of hits), D-prime score, and RT with n-back load (2, 3, 4-back), emotional condition (control, stress) as within subject, and stimulation groups as between-subject factors. Secondary rmANOVAs were conducted for each WM load condition with emotional condition as within-subject and stimulation group as between-subject factors. For EEG analyses, time-averaged P200 and P300amplitudes, and time– and frequency-averaged theta ERS and alpha ERD activity were analyzed using rmANOVAs with emotional condition as within-subject, and stimulation group as between-subject factor for each WM load (2, 3, 4-Back). For side-effects during and after stimulation, rmANOVAs were conducted with emotional condition as within-subject, and stimulation group as between-subject factor. For assessing proper blinding, chi-square tests were conducted. A one-way ANOVA for age, chronotype (MEQ scores), and state anxiety (STAI-S) were performed with stimulation group as a between-subject factor.

Sphericity of the data was tested for all ANOVAs with the Mauchly test, and Greenhouse-Geisser corrections were applied when appropriate. For post-hoc tests, which were conducted in case of significant ANOVA results, Fisher’s Least Significant Difference (LSD) test was used. The critical alpha level was set at 0.05 for all tests. Correlations were performed using Pearson’s correlation for WM performance (Hits and D-Prime) with EEG (Alpha ERD, Theta ERS, the P200 amplitude, and the P300 amplitude) and physiological markers (cortisol levels before and after task performance, HR, and HRV during the task). EEG data were plotted using MATLAB (R2020b) and the open-source EEG data analysis software EEGLAB (55).

## Acknowledgments

This work was funded by DAAD (Deutscher Akademischer Austauschdienst) GSSP (Graduate School Scholarship Programme) grant to S.R.

## Author Contributions

S.R., Y.F., M.M.S., and M.A.N. designed the experiment. S.R. collected the data. S.R. analyzed behavior, physiological, and EEG data. M.C. analyzed cortisol and immunological data. F.Y. did the modeling for the stimulation montage. S.R. prepared the figures. S.R. wrote the main manuscript text. T.K. and M.A.N. critically revised the manuscript and provided expert comments.

## Classification

Biological Science – Neuroscience; Social Science – Psychological and Cognitive Sciences

