## Supplementary Material for "Modulating Prefrontal Cortex Activity to Alleviate Stress-Induced Working Memory Deficits: A Transcranial Direct Current (tDCS) Study"

**1. Supplementary Methods –**

**1.1 Heart Rate (HR) and Heart Rate Variability (HRV) Data –** HR was continuously monitored during the experiment using two bipolar electrodes connected to an EEG jack box (Neurone, Finland)— one electrode was attached below the right clavicle and the other electrode was attached above and left to the umbilicus. HR data were sampled at 2000 Hz and analyzed using HEPLAB (1), an EEGLAB (2) plugin, along with custom MATLAB scripts. Inter-beat intervals (IBIs) were calculated, and HR was computed for different time points in the experiment. HRV was determined using the Root Mean Square of Successive Differences (RMSSD) method (3) via custom MATLAB scripts. HR and HRV data were baseline-corrected to the first time point of the experiment. One participant's data (from the sham group) was rejected due to bad data quality from which IBIs were not recognizable.

**1.2 Cortisol –** Saliva samples were collected at various time points during the experiment using Salivettes (Sarstedt, Nümbrecht, Germany), Cortisol levels were analyzed using a Cortisol saliva ELISA (TECAN/IBL International, Männedorf, Switzerland) following manufacturer instructions. Absorbance at 450nm was measured using a GloMax® Multimode Microplate reader system (Promega, Madison, United States).

**1.3 Cytokines –** Cytokine levels in saliva were measured twice per sample using the LEGENDplex Human Essential Immune Response panel (BioLegend, San Diego, United States), as described in previous studies and in our previous work (4). Cytokines targeted for analysis were Interferon-gamma (IFN-γ), Interleukin – 1beta (IL-1β), Interleukin – 6 (IL-6), Interleukin – 4 (IL-4), Interleukin – 8 (IL-8), and Tumor necrosis factor alpha (TNFα). We selected this profile of six essential cytokines to obtain a broad overview of stress effects on cytokines by targeting a mix of pro-inflammatory (IFN-γ, IL8, IL-1β, TNFα, and IL-6) and anti-inflammatory (IL-4) cytokines, in accordance with previous studies (5, 6). Cytokine data from only 70 participants were analyzed due to limited availability of kits and insufficient saliva samples from some participants.

**1.4 EEG data Acquisition and processing –** Data preprocessing was uniform for all datasets, and a pipeline described in our previous study was used (4). The data were epoched into 2-second segments with no baseline for resting-state data and into -1 to 2s blocks relative to stimulus presentation in the task-based analysis. The continuous n-back data were divided into three groups based on n-back levels. The task-based analysis was conducted for correctly recognized epochs (hits + correct rejections) which were baseline-corrected from -250 to -50 ms relative to stimulus presentation. Epochs exceeding amplitudes of ±100 μV were rejected resulting in an average number of epochs over all subjects of 122±26 for 2-Back, 114±24 for 3-back, and 103±22 for 4-Back load conditions. We also conducted a visual check of data quality, and noisy channels were rejected. We then ran an ICA decomposition using the EEGLAB 'runICA' command and the resulting decompositions were labeled using the IClabel plugin (7). Decompositions having more than 70% probability of being an eye, muscle, line, or noise artifact were rejected from the data. The data of three participants (one from each group) were removed due to technical problems during recording. For analysing concurrent tDCS-EEG data, we followed technical guidelines described in previous work (8).

**1.5 Power Analysis –** The power analysis employed the EEGLAB 'STUDY' function and custom MATLAB scripts. The *spectopo* function was used for power estimation with a window size of 998ms and an overlap of 495ms, and spectra were generated for the electrodes of interest based on the frequency band. We analyzed theta power in the frequency range between 4 to 8 Hz from frontal electrodes (Fz, F1, F2) and alpha power in the frequency range between 8 to 13 Hz from parietal electrodes (Pz, P1, P2) for resting state data for eyes open (EO) and eyes closed (EC) conditions separately.

**1.6 Time frequency (TF) Cluster analysis –** We averaged the data from frontal (Fz, F1, and F2) and parietal (Pz, P1, and P2) channels for task-based theta and alpha activity respectively, and did Montecarlo-based permutation statistics with cluster correction on the time x frequency data (Fieldtrip toolbox in EEGLAB (2, 9)) to identify the relevant time range for comparisons between conditions. We performed 8000 random permutations to establish a distribution of cluster statistics to which each cluster t-value was compared. A threshold of alpha </= 0.05 (two-tailed) was applied to this random distribution to identify significant clusters. For TF data, these were identified based on their spectral and time adjacency. Analysis was done for each WM load (2, 3, and 4-back) separately as shown in the supplementary material.

**1.7 Data Analysis -** Repeated Measure analyses of variance (rmANOVA) were performed for subjective data (positive, negative affect scores, and STAI-S scores), with emotional condition (control, stress), and time (before and after WM task performance) as within-subject factors, and stimulation group (sham, dlPFC, and vmPFC) as the between-subject factor. rmANOVAs for cytokines were performed with emotional condition (control, stress) and time (before, immediately after task performance, and after a 40-minute break) as within-subject factors, and stimulation group as between-subject factor. Similar rmANOVAs were also conducted for HRV with emotional condition and time (t1-t23) as within-subject, and stimulation group as between-subject factors. The first time point of the experiment before stress induction, as described in previous work (4), was taken as baseline, and then the change from baseline after emotional/control clip presentation to the end of the experiment was analyzed for HRV data. rmANOVAs were conducted for reaction time (RT) with n-back load (2, 3, 4-back), emotional condition (control, stress) as within-subject, and stimulation groups as between-subject factors. Secondary rmANOVAs were conducted for each WM load condition with emotional condition as within-subject and stimulation group as between-subject factors. For EEG analyses, time-averaged P200 and P300 amplitudes, and time- and frequency-averaged theta ERS and alpha ERD activity were analyzed using rmANOVAs with emotional condition as within-subject, and stimulation group as between-subject factor for each WM load (2, 3, 4-Back) (2-back and 4-back data shown here). Averaged resting state EEG power data for the frequency of interest and EO and EC conditions were statistically analyzed via rmANOVAs with emotional condition (control, stress), resting state blocks (before and after WM performance) as within-subject factors, and stimulation group as between subject factor. For side-effects during and after stimulation, rmANOVAs were conducted with emotional condition as within-subject, and stimulation group as between-subject factor. One-way ANOVAs for age, chronotype (MEQ scores), and state anxiety (STAI-S) were performed with the stimulation group as a between-subject factor.

### 2. Supplementary Results

**2.1 Descriptive Statistics –** For demographic variables, we did not find any group differences with respect to age, MEQ, or STAI – trait anxiety scores between participants (see Table S1).

*Supplementary Table 1: Baseline values of Age, MEQ, and STAI-Trait values of the experimental groups. Mean values with SD are shown. $\eta_p 2$= partial eta squared, MEQ = morningness-eveningness questionnaire, STAI-T = state-trait anxiety inventory (trait).*

| | Sham (n = 26) | dlPFC (n = 26) | vmPFC (n = 26) | Statistics |
|---|---|---|---|---|
| **Age** | 26 ± 3.69 | 24.27 ± 3.54 | 25.08 ± 4.97 | ([F (2, 75) = 1.149, p = 0.323, ηp2 = 0.03]) |
| **MEQ** | 47.19(9.73) | 46.64(9.58) | 49.19(8.63) | ([F (2, 74) = .589, p = 0.589, ηp2 = 0.014]) |
| **STAI-T** | 39.4(6.78) | 38.52(9.42) | 41.08(9.76) | [F (2, 73) = .561, p = 0.573, ηp2 = 0.015] |

**2.2 – Side Effects –** We found a significant effect of simulation group on pain perceived during stimulation, with the vmPFC group reporting less pain compared to the dlPFC (p = 0.037) and sham groups (p = 0.003). We also found a significant effect of emotional condition on nervousness, where the stress condition resulted in increased nervousness as compared to the control group (p = 0.044) (see Table S3).

*Supplementary Table 2 Mean scores of self-report measures of tDCS for the experimental groups. Mean values with SD are shown for the groups.*

| | Sham Control | Sham Stress | dlPFC Control | dlPFC Stress | vmPFC Control | vmPFC Stress |
|---|---|---|---|---|---|---|
| **Itching** | 1.65 ± 1.67 | 1.7 ± 1.64 | 1.26 ± 1.4 | 1.26 ± 1.4 | 1.2 ± 1.4 | 1.57 ± 1.52 |
| **Tingling** | 2.11 ± 1.65 | 1.69 ± 1.66 | 1.15 ± 1.51 | 1.42 ± 1.6 | 1.15 ± 1.43 | 1.57 ± 1.27 |
| **Burning** | 1.92 ± 1.62 | 1.88 ± 1.7 | 1.42 ± 1.55 | 1.23 ± 1.36 | 1.11 ± 1.3 | 1.19 ± 1.2 |
| **Pain** | 1.11 ± 1.27 | 1.23 ± 1.45 | .8 ± 1.2 | 1 ± 1.54 | .34 ± .68 | .23 ± .51 |
| **Redness** | .42 ± .75 | .76 ± 1.14 | .73 ± 1.07 | .88 ± 1.24 | 1.03 ± 1.21 | .65 ± .97 |
| **Headache** | .19 ± .49 | .3 ± .88 | .03 ± 1.96 | .46 ± 1.02 | .3 ± .83 | .15 ± .37 |
| **Fatigue** | .38 ± .89 | .27 ± .53 | .38 ± .94 | .61 ± 1.09 | .42 ± .94 | .23 ± .58 |
| **Concentration** | .3 ± .83 | .57 ± 1.17 | .57 ± 1.1 | 1.23 ± 1.79 | .46 ± .94 | .26 ± .6 |
| **Nervousness** | .15 ± .61 | .34 ± .74 | .15 ± .46 | .61 ± 1.2 | .11 ± .43 | .11 ± .43 |

*Supplementary Table 3 Main and interaction effects of ANOVAs conducted for self-reported scores of side effects during and after stimulation. Significant p-values are marked in bold. d.f. = degrees of freedom, $\eta_p 2$= partial eta squared.*

| | | Factors | d.f., | F Value | P - value | $\eta_p^2$ |
|---|---|---|---|---|---|---|
| **During Stimulation** | **Itching** | **Emotional Condition** | 1,2 | .633 | .429 | .008 |
| | | **Stimulation group** | 2,75 | .569 | .569 | .015 |
| | | **Emotional condition x Stimulation group** | 2,75 | .251 | .779 | .007 |
| | **Tingling** | **Emotional Condition** | 1,2 | .196 | .659 | .003 |
| | | **Stimulation group** | 2,75 | 1.84 | .165 | .047 |

| | | | | | | |
|---|---|---|---|---|---|---|
| | | **Emotional condition x Stimulation group** | 2,75 | 1.65 | .199 | .042 |
| | **Burning** | **Emotional Condition** | 1,2 | .108 | .744 | .001 |
| | | **Stimulation group** | 2,75 | 2.38 | .1 | .06 |
| | | **Emotional condition x Stimulation group** | 2,75 | .249 | .78 | .007 |
| | **Pain** | **Emotional Condition** | 1,2 | .242 | .624 | .003 |
| | | **Stimulation group** | 2,75 | 4.88 | **.01** | .115 |
| | | **Emotional condition x Stimulation group** | 2,75 | .503 | .607 | .013 |
| **After Stimulation** | **Redness** | **Emotional Condition** | 1,2 | .082 | .775 | .001 |
| | | **Stimulation group** | 2,75 | .575 | .565 | .015 |
| | | **Emotional condition x Stimulation group** | 2,75 | 2.65 | .077 | 0.06 |
| | **Headache** | **Emotional Condition** | 1,2 | 1.4 | .239 | .018 |
| | | **Stimulation group** | 2,75 | .012 | .988 | .0 |
| | | **Emotional condition x Stimulation group** | 2,75 | 2.37 | .1 | 0.06 |
| | **Fatigue** | **Emotional Condition** | 1,2 | .055 | .814 | .001 |
| | | **Stimulation group** | 2,75 | .513 | .601 | .014 |
| | | **Emotional condition x Stimulation group** | 2,75 | 1.43 | .246 | .037 |
| | **Concentration** | **Emotional Condition** | 1,2 | 2.1 | .151 | .027 |
| | | **Stimulation group** | 2,75 | 2.9 | .058 | .073 |
| | | **Emotional condition x Stimulation group** | 2,75 | 2.12 | .12 | .054 |
| | **Nervousness** | **Emotional Condition** | 1,2 | 4.18 | **.044** | .053 |
| | | **Stimulation group** | 2,75 | 1.73 | .214 | .044 |
| | | **Emotional condition x Stimulation group** | 2,75 | 1.57 | .214 | .04 |

**2.3 Subjective Measures of Stress –** We analyzed PANAS and STAI to assess subjective measures of stress and if stimulation changed subjective feelings of stress. The rmANOVA conducted for positive affect, and negative affect obtained by the PANAS, and the STAI-S scores revealed a common pattern of increased state anxiety, negative affect, and decreased positive affect scores in the stress as compared to the control condition. No differences related to stimulation were found. Please refer to Table S5 for ANOVA results and to Figure S1.

*Supplementary Table 4 Mean scores of questionnaire data (positive, negative affect, and STAI-S scores) before and after the working memory (WM) task for the experimental groups. Mean values with SD are shown for the groups.*

| | | **Sham Control** | **Sham Stress** | **dlPFC Control** | **dlPFC Stress** | **vmPFC Control** | **vmPFC Stress** |
|---|---|---|---|---|---|---|---|
| **Positive Affect** | **Before WM task** | 28.23±8.6 | 23.11±6.6 | 27.23±9.8 | 24.07±5.7 | 27.46±9.2 | 23.92±6.9 |
| | **After WM task** | 28.38±8.6 | 26.19±6.6 | 29.1±10.6 | 25.3±9.6 | 28.11±9.7 | 26.6±10.1 |
| **Negative Affect** | **Before WM task** | 13.42±3.9 | 24.88±7.2 | 13.61±4.9 | 25.42±9.7 | 13.04±3.1 | 23.92±8.9 |
| | **After WM task** | 14.26±4.1 | 16.88±7.1 | 14±4.8 | 16.26±6.9 | 12.9±4.2 | 15.53±5.3 |
| **State Anxiety** | **Before WM task** | 36.38±5.6 | 51.07±9.5 | 35.1±5.1 | 51.53±10.2 | 35.96±6 | 52.15±10.7 |
| | **After WM task** | 37.46±5.9 | 41.07±8.8 | 35.96±5.2 | 43±9.3 | 38.08±6.4 | 42.03±9.05 |

*Supplementary Table 5 Main and interaction effects of the ANOVAs conducted for questionnaire data (positive, negative affect, and STAI-S scores). Significant p-values are* *marked in bold. d.f. = degrees of freedom, $\eta_p2$= partial eta squared, STAI-S = state-trait anxiety inventory - state.*

| | Factors | d.f., Error | F Value | P - value | $\eta_p^2$ |
|---|---|---|---|---|---|
| **Positive Affect Scores** | **Emotional Condition** | 1, 75 | 19.226 | **<0.001** | 0.204 |
| | **Time** | 1, 75 | 7.2 | **0.009** | 0.088 |
| | **Stimulation Group** | 2, 75 | 0.002 | 0.998 | 0.000 |
| | **Emotional Condition x Time** | 1, 75 | 3.221 | 0.077 | 0.041 |
| | **Emotional condition x Stimulation Group** | 2, 75 | 0.238 | 0.788 | 0.006 |
| | **Stimulation group x Time** | 2, 75 | 0.005 | 0.995 | 0.000 |
| | **Emotional Condition x Time x Stimulation** | 2, 75 | 1.716 | 0.197 | 0.044 |
| **Negative Affect Scores** | **Emotional Condition** | 1, 75 | 114.56 | **<0.001** | 0.604 |
| | **Time** | 1, 75 | 99.86 | **<0.001** | 0.571 |
| | **Stimulation Group** | 2, 75 | 0.354 | 0.703 | 0.009 |
| | **Emotional Condition x Time** | 1, 75 | 103.2 | **<0.001** | 0.579 |
| | **Emotional condition x Stimulation Group** | 2, 75 | 0.025 | 0.975 | 0.001 |
| | **Stimulation group x Time** | 2, 75 | 0.371 | 0.692 | 0.01 |
| | **Emotional Condition x Time x Stimulation** | 2, 75 | 0.165 | 0.849 | 0.004 |
| **STAI - S** | **Emotional Condition** | 1, 75 | 122.82 | **<0.001** | 0.621 |
| | **Time** | 1, 75 | 55.33 | **<0.001** | 0.425 |
| | **Stimulation Group** | 2, 75 | 0.089 | 0.915 | 0.002 |
| | **Emotional Condition x Time** | 1, 75 | 125.47 | **<0.001** | 0.626 |
| | **Emotional condition x Stimulation Group** | 2, 75 | 0.653 | 0.524 | 0.017 |
| | **Stimulation group x Time** | 2, 75 | 0.110 | 0.896 | 0.003 |
| | **Emotional Condition x Time x Stimulation** | 2, 75 | 0.705 | 0.497 | 0.018 |

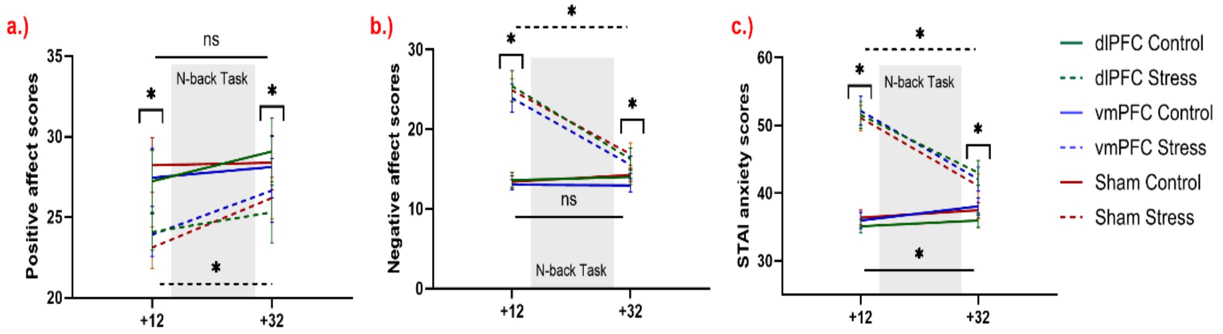


*Supplementary Figure 1 Subjective measures of affect before and after WM performance. a.) Positive affect scores before and after WM task conduction combined with tDCS for different emotional condition and stimulation groups are shown. b.) Negative affect scores before and after task conduction combined with tDCS are presented. c.) STAI-S anxiety scores before and after task conduction combined with stimulation are shown. For these subjective scores, the y-axis denotes the scores in the respective questionnaires and the x-axis shows the time course relative to stressor onset. Post-hoc tests conducted based on the significant emotional x time interaction in the respective ANOVAs showed increased state anxiety, increased negative affect, and decreased positive affect scores*

*in the stress condition as compared to the control conditions before the start of the WM task and after task performance as denoted by asterisks (*). We also found a significant decrease in state anxiety, negative affect and increase in positive affect after compared to before task performance in the stress condition for all stimulation groups as shown by a horizontal line marked by an asterisk (*), and a significant increase of state anxiety after task performance in the control condition as shown by a horizontal line marked by an asterisk (*). Error bars show ± SEM.*

**2.4 – Cytokines:** The rmANOVA performed for cytokine analysis revealed a significant main effect of time for all cytokines except for IL-8. For IL-8, a significant interaction effect of emotional condition, time, and stimulation group was observed [F (3.47, 119.7) = 2.73, p = .039]. The post hoc tests revealed increased IL-8 levels in the stress as compared to the control condition after the 40 mins break after WM task performance only in the dlPFC group (p = 0.03), while no such significant differences were found for other time points or in other groups. For IL-1, we observed a significant main effect of emotional condition [F (1, 69) = 5.118, p = .027] revealing larger IL-1 in the control compared to the stress condition (p = 0.027).

*Supplementary Table 6 Main and interaction effects of the rmANOVAs conducted for cytokine (IL-6, IL-4, TNF-α, IFN- γ, IL-1, and IL-8) levels. Significant p-values are marked in bold. d.f. = degrees of freedom, $\eta_p 2$= partial eta squared.*

| | **Factors** | **d.f., Error** | **F Value** | **P - value** | **$\eta_p^2$** |
|---|---|---|---|---|---|
| **Interleukin-6 (IL-6)** | **Emotional Condition** | 1 (69) | 2.13 | .149 | 0.03 |
| | **Time** | 2 (138) | 11.775 | **<.001** | 0.146 |
| | **Stimulation Group** | 2 (69) | .191 | .826 | 0.006 |
| | **Emotional Condition x Time** | 2 (138) | 2.403 | .094 | 0.034 |
| | **Emotional condition x Stimulation Group** | 2(69) | .416 | .661 | 0.012 |
| | **Stimulation group x Time** | 4 (138) | .488 | .744 | 0.014 |
| | **Emotional Condition x Time x Stimulation Group** | 4 (138) | .57 | .685 | 0.015 |
| **Interleukin -4 (IL-4)** | **Emotional Condition** | 1 (69) | .180 | .672 | 0.003 |
| | **Time** | 2 (138) | 23.06 | **<.001** | 0.251 |
| | **Stimulation Group** | 2 (69) | 1.060 | .352 | 0.03 |
| | **Emotional Condition x Time** | 1.73 (119.85) | 2.533 | .083 | 0.035 |
| | **Emotional condition x Stimulation Group** | 2(69) | .389 | .679 | 0.011 |
| | **Stimulation group x Time** | 4 (138) | .602 | .662 | 0.017 |
| | **Emotional Condition x Time x Stimulation Group** | 3.47 (119.85) | 1.146 | .336 | 0.032 |

| | | | | | |
|---|---|---|---|---|---|
| **Tumor Necrosis Factor - Alpha TNF-α** | **Emotional Condition** | 1 (69) | .743 | .392 | 0.011 |
| | **Time** | 1.73 (119.33) | 17.447 | **<.001** | 0.202 |
| | **Stimulation Group** | 2 (69) | .680 | .510 | 0.019 |
| | **Emotional Condition x Time** | 2 (138) | 1.592 | .207 | 0.023 |
| | **Emotional condition x Stimulation Group** | 2(69) | .382 | .684 | 0.011 |
| | **Stimulation group x Time** | 3.46 (119.33) | .358 | .811 | 0.01 |
| | **Emotional Condition x Time x Stimulation Group** | 4 (138) | 1.837 | .125 | 0.051 |
| **Interferon – Gamma (IFN- γ)** | **Emotional Condition** | 1 (69) | 1.195 | .278 | 0.017 |
| | **Time** | 2 (138) | 31.037 | **<.001** | 0.184 |
| | **Stimulation Group** | 2 (69) | .716 | .492 | 0.02 |
| | **Emotional Condition x Time** | 2 (138) | 1.920 | .151 | 0.027 |
| | **Emotional condition x Stimulation Group** | 2(69) | .564 | .571 | 0.016 |
| | **Stimulation group x Time** | 4(138) | .454 | .770 | 0.013 |
| | **Emotional Condition x Time x Stimulation Group** | 4(138) | 1.277 | .282 | 0.036 |
| **Interleukin -1 (IL-1)** | **Emotional Condition** | 1, 69 | 5.118 | **.027** | 0.069 |
| | **Time** | 1.8, 124.6 | 28.855 | **<.001** | 0.173 |
| | **Stimulation Group** | 2, 69 | 1.085 | .344 | 0.03 |
| | **Emotional Condition x Time** | 2, 138 | .553 | .577 | 0.008 |
| | **Emotional condition x Stimulation Group** | 2, 69 | .997 | .374 | 0.028 |
| | **Stimulation group x Time** | 3.6, 124.6 | 1.273 | .284 | 0.036 |
| | **Emotional Condition x Time x Stimulation Group** | 4, 138 | 2.33 | .059 | 0.063 |
| **Interleukin -8 (IL-8)** | **Emotional Condition** | 1, 69 | .019 | .890 | 0.000 |
| | **Time** | 2.4, 82.7 | 1.041 | .324 | 0.015 |
| | **Stimulation Group** | 2, 69 | .256 | .775 | 0.007 |
| | **Emotional Condition x Time** | 1.73, 119.7 | .544 | .557 | 0.008 |
| | **Emotional condition x Stimulation Group** | 2, 69 | 2.938 | .06 | 0.078 |
| | **Stimulation group x Time** | 2.4, 82.69 | .1.367 | .261 | 0.038 |
| | **Emotional Condition x Time x Stimulation Group** | 3.47, 119.7 | 2.731 | **.039** | 0.073 |

***HRV:*** In the HRV analysis, the rmANOVA showed only a significant main effect of time [F (5.5, 405.6) = 11.3, p<.001], but no significant effects were observed for emotional conditions, stimulation groups or respective interactions (see figure S2).

*Supplementary Table 7 Main and interaction effects of the rmANOVAs conducted for HRV levels. Significant p-values are marked in bold. d.f. = degrees of freedom, $\eta_p 2$= partial eta squared.*

| | **Factors** | **d.f., Error** | **F Value** | **P - value** | **$\eta_p^2$** |
|---|---|---|---|---|---|
| **HRV** | **Emotional Condition** | 1,74 | 1.454 | 0.232 | 0.019 |
| | **Time** | 5.5,405.6 | 11.3 | **<0.001** | 0.133 |
| | **Stimulation Group** | 2,74 | 0.53 | 0.592 | 0.014 |

| | | | | |
|---|---|---|---|---|
| **Emotional Condition x Time** | 8.07, 597.1 | 0.718 | 0.677 | 0.01 |
| **Emotional condition x Stimulation Group** | 2, 74 | 1.25 | 0.293 | 0.033 |
| **Stimulation group x Time** | 10.96, 405.6 | 0.87 | 0.569 | 0.023 |
| **Emotional Condition x Time x Stimulation Group** | 16.14, 597.1 | 0.934 | 0.53 | 0.025 |

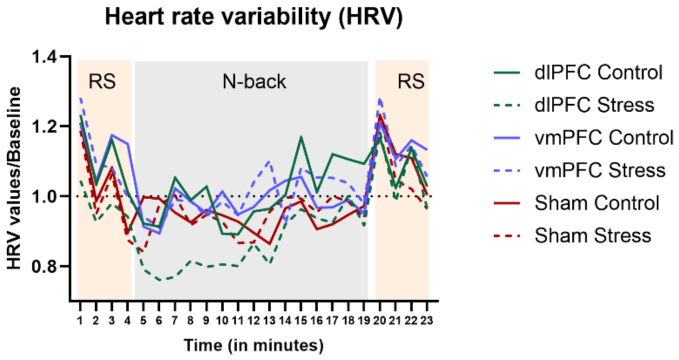


*Supplementary Figure 2 Heart rate variability (HRV) before, during, and after task performance. HRV was calculated based on RMSSD measures. The y-axis denotes HR changes from baseline (first time point of the experiment as decribed in the methods section), and the x-axis denotes each minute of the experiment (from the end of the stress induction procedure). Resting state (RS) and n-back task time frames are highlighted in the background with grey and orange colored watermarks (orange represents the resting state and grey the n-back task).*

**2.5 Reaction Time (RT):** We performed first a 3-way rmANOVA for all loads combined and then a 2-way rmANOVA for each WM load for RT. See Table S7 for the ANOVA results and Fig S2 for data.

The 3-way rmANOVA showed significant main effects of WM load [F (1.7, 128.2) = 38.37, p <.001]. The respective post hoc tests conducted for WM load showed lower RT in the 2-back compared to the 3-back and 4-back load (all p <.001), and lower RT in the 3-back compared to the 4-back load (p=.009). No other significant main or interaction effects were seen of emotional condition or stimulation.

Secondary ANOVAs conducted for each n-back levels did not reveal any effects of emotional condition or stimulation in any of the n-back loads.

*Supplementary Table 8 Main and interaction effects of the rmANOVAs conducted for RT data of the WM task. First, a 3-way RM ANOVA including all loads was performed and* *then separate ANOVAs for each load were performed. Significant p-values are marked in bold. d.f. = degrees of freedom, $\eta_p$2= partial eta squared.*

| | | Factors | d.f., Error | F Value | P - value | $\eta_p^2$ |
|---|---|---|---|---|---|---|
| **Reaction Time** | **All loads** | **Emotional Condition** | 1, 72 | 0.545 | 0.463 | 0.008 |
| | | **Load** | 1.78, 128.6 | 38.37 | **<.001** | 0.348 |
| | | **Stimulation Group** | 2, 72 | 0.054 | 0.948 | 0.001 |
| | | **Emotional Condition x Load** | 1.85, 132.9 | 0.58 | 0.55 | 0.008 |
| | | **Emotional condition x Stimulation Group** | 2, 72 | 2.031 | 0.139 | 0.053 |
| | | **Stimulation group x Load** | 3.57, 128.6 | 1.38 | 0.244 | 0.037 |
| | | **Emotional Condition x Load x Stimulation Group** | 3.69, 132.9 | 0.385 | 0.819 | 0.011 |
| | **2-Back** | **Emotional Condition** | 1, 72 | 0.921 | 0.34 | 0.013 |
| | | **Stimulation Group** | 2, 72 | 0.976 | 0.382 | 0.026 |
| | | **Emotional condition x Stimulation Group** | 2, 72 | 0.133 | 0.876 | 0.004 |
| | **3-Back** | **Emotional Condition** | 1, 72 | 0.146 | 0.704 | 0.002 |
| | | **Stimulation Group** | 2, 72 | 0.029 | 0.971 | 0.001 |
| | | **Emotional condition x Stimulation Group** | 2, 72 | 1.3 | 0.279 | 0.035 |
| | **4-Back** | **Emotional Condition** | 1, 72 | 0.657 | 0.42 | 0.009 |
| | | **Stimulation Group** | 2, 72 | 0.096 | 0.9 | 0.003 |
| | | **Emotional condition x Stimulation Group** | 2, 72 | 1.27 | 0.287 | 0.034 |

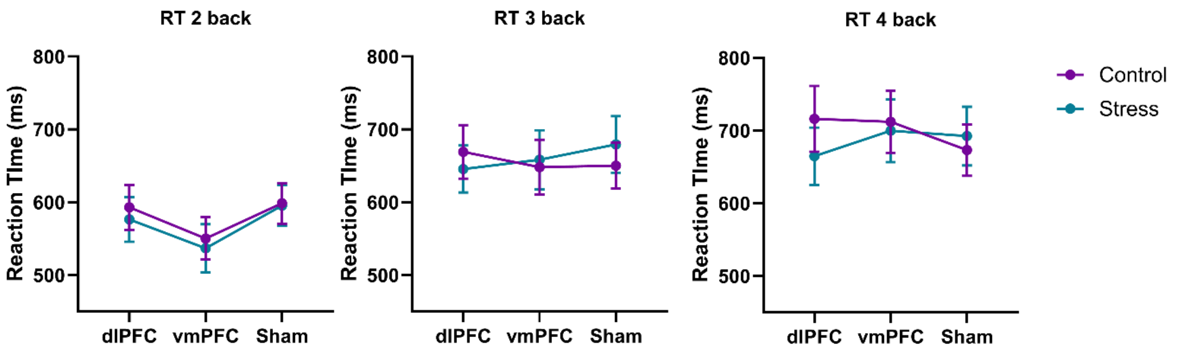


*Supplementary Figure 2 Line Plots to show reaction time (RT) for hits for all three n-back load conditions. The y-axis shows RT in milliseconds (ms) and the x-axis shows different stimulation groups. Conditions (control, stress) are indicated by the color of the lines. Error bars show ± SEM.*

**2.6 EEG Results (2-Back):** We analyzed the averaged P200 ERP component, P300 ERP component, task-based theta ERS, and task-based alpha ERD values (see supplementary Fig S3, Table S8).

**2.6.1 P200 ERP Component:** We found no significant main or interaction effects of emotional condition or stimulation group with respect to the P200 ERP component.

**2.6.2 P300 ERP Component:** We found a significant main effect of emotional condition on the P300 ERP amplitude [$F(1, 72) = 6.85$, $p = .011$] but no significant main effect of stimulation group or interaction between these factors..

**2.6.3 Theta ERS:** We found no significant main or interaction effect of emotional condition and stimulation group on task-based theta ERS activity.

**2.6.4 Alpha ERD:** We found no significant main or interaction effect of emotional condition and stimulation group on task-based alpha ERD activity.

*Supplementary Table 9 Main and interaction effects of the ANOVAs conducted for EEG indexes in the 2-Back load WM task condition (P200 component, P300 component, Theta ERS, and Alpha ERD values). Significant p-values are marked in bold. d.f. = degrees of freedom, $\eta_p 2$= partial eta squared.*

| | **Factors** | **d.f., Error** | **F Value** | **P - value** | **$\eta_p^2$** |
|---|---|---|---|---|---|
| **P200** | **Emotional Condition** | 1,72 | 2.37 | .128 | .032 |
| **ERP (Cz)** | **Stimulation Group** | 2,72 | 1.08 | .344 | .029 |
| | **Emotional Condition x Stimulation Group** | 2,72 | .6 | .55 | .017 |
| **P300** | **Emotional Condition** | 1,72 | 6.85 | **.011** | .087 |
| **ERP (Pz)** | **Stimulation Group** | 2,72 | .055 | .946 | .002 |
| | **Emotional Condition x Stimulation Group** | 2,72 | .515 | .6 | .014 |
| **Theta** | **Emotional Condition** | 1,72 | .024 | .877 | .0 |
| **ERS** | **Stimulation Group** | 2,72 | 1.99 | .147 | .052 |
| | **Emotional Condition x Stimulation Group** | 2,72 | 1.07 | .349 | .029 |
| **Alpha** | **Emotional Condition** | 1,72 | 1.01 | .32 | .014 |
| **ERD** | **Stimulation Group** | 2,72 | 1.45 | .242 | .039 |
| | **Emotional Condition x Stimulation Group** | 2,72 | .058 | .943 | .002 |

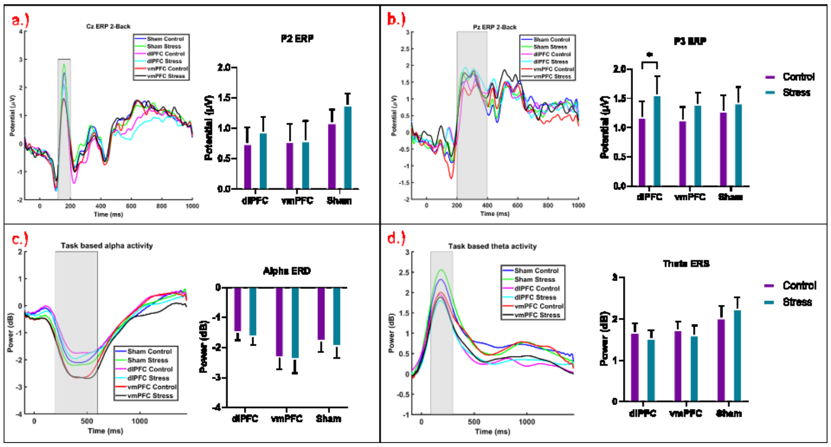


*Supplementary Figure 4 EEG results for the 2-Back load. a.) averaged ERP for all groups and conditions for EEG channel Cz. The x-axis represents time (in ms) and the y-axis shows the ERP amplitude (µV). The P200 (120-200ms) component is highlighted by a gray box from which the averaged amplitude values were used for statistical analysis shown next to it in the bar plot. The bar plot shows the averaged P200 component values for all groups and conditions with the x-axis showing groups and the y-axis the potential amplitude (µV). b.) averaged ERP for all groups and conditions for the EEG channel Pz to show the P300 (200-400ms) component amplitude plotted as for P2. c.) averaged task-based alpha activity from parietal channels (Pz, P1, and P2) for all groups and conditions with the x-axis representing time (ms) and the y-axis power (in dB). The time range of interest for ERD (200-600 ms) is highlighted in grey. ERD averaged over this time range are plotted in a bar graph with the x-axis showing groups and the y-axis power (dB). d.) averaged task-based theta activity in frontal channels (Fz, F1, and F2) for ERS (80-300ms) plotted similarly as for alpha ERD. Error bars denote ± SEM.*

**2.7: TF Cluster Results (2-back):**

We analyzed channel-averaged task-based theta and alpha activity over the whole 2-back epoch between conditions using Montecarlo-based permutation statistics with cluster correction

**2.7.1 Theta Activity:** No significant difference clusters were found between emotional conditions or stimulation groups in the 2-back load WM task for theta activity (see supplementary Figure S4a).

**2.7.2 Alpha Activity:** No significant difference clusters were found between emotional conditions or stimulation groups in the 2-back load WM task for alpha activity (see supplementary Figure S4b).

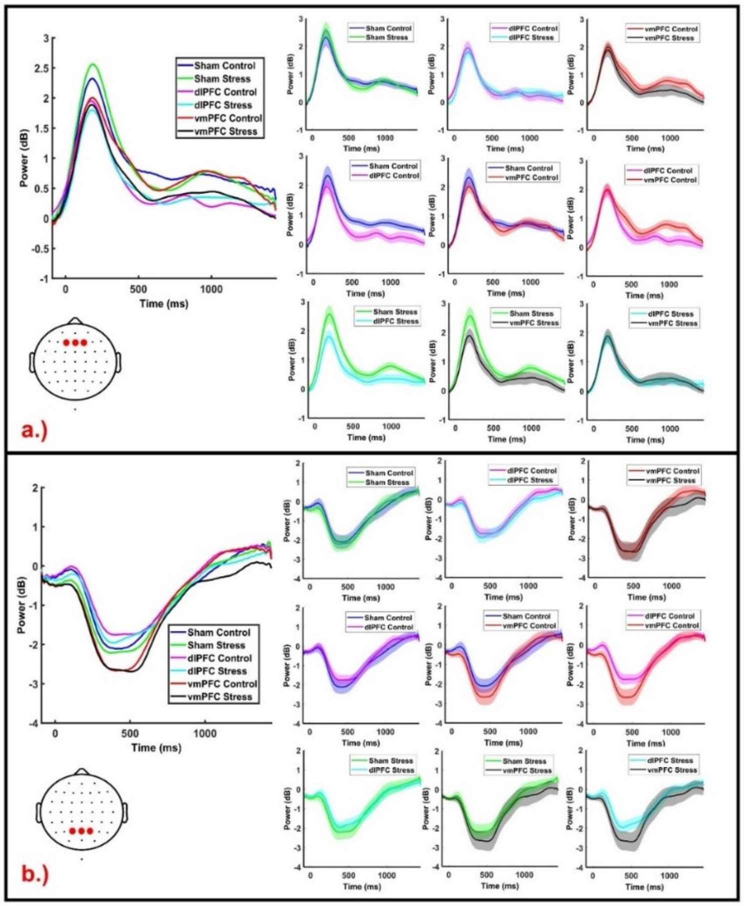


*Supplementary Figure 5 shows time-frequency plots for task-related theta and alpha activity during for 2-Back task performance. a.) Theta ERD/ERS activity derived from frontal electrodes (Fz, F1, and F2) are shown with power on the y-axis and time (in ms) on the x-axis in the left figure with averaged theta (4-8 Hz) activity over the time course for different stimulation and emotional conditions plotted together. The cartoon head below shows the positions of the target electrodes. The right side of the figures shows difference plots between conditions, as indicated in the legend, with the shaded colored zones being the standard deviation for each condition. Cluster-based permutation testing was applied to identify significant clusters of time bins in the analysis. b.) Alpha ERD/ERS activity from parietal electrodes (Pz, P1, and P2) are shown with power indicated by the y-axis and time (in ms) by the x-axis, plotted as described above.*

**2.8 TF Cluster Results (3-Back):** We analyzed channel-averaged task-based theta and alpha activity over all 3-back epochs between conditions using Montecarlo-based permutation statistics with cluster correction, as described for the 2-back load condition. Refer to Table S9 for cluster and time range details.

**2.8.1 Theta Activity:** We found reduced theta activity during late processing of stimuli similar to a previous study using similar clips (10) in the stress group as compared to the control group in the sham stimulation condition, with a positive cluster in the time range of 910 ms to 1366 ms from stimulus onset. No other significant cluster was seen for other groups or conditions (see supplementary Figure S5a).

**2.8.2 Alpha Activity:** We found significant differences between the vmPFC and dlPFC stimulation groups in both stress and control conditions. Specifically, in the control condition we found a positive cluster in the time range of 182 – 598 ms, and in the stress condition, we found a positive cluster in the time range of 154 – 628 ms with a larger alpha ERD in the vmPFC group as compared to the dlPFC group (see supplementary Figure S5b).

*Supplementary Table 10 Cluster information for the TF analysis of the 3-back load for task-based theta and alpha activity. The number of clusters, cluster t-values, p-values, and time range in each cluster are given for each analysis. The sign of the t value denotes the direction of effects, e.g. for condition 1 vs condition 2 for each analysis, a positive cluster shows a higher amplitude in condition 1, and a negative cluster shows a larger amplitude in condition 2. Only significant clusters are shown. Cluster t-values were calculated using cluster-based permutation testing using the Fieldtrip toolbox in EEGLAB.*

| | Condition 1 | Condition 2 | Number of Clusters | t-value | P-value | Time range |
|---|---|---|---|---|---|---|
| **Theta Activity** | **Sham Control** | **Sham Stress** | **1** | **124.62** | **0.0085** | **910 – 1366** |
| | **dlPFC Control** | **dlPFC Stress** | 0 | | | |
| | **vmPFC Control** | **vmPFC Stress** | 0 | | | |
| | **Sham Control** | **dlPFC Control** | 0 | | | |
| | **Sham Control** | **vmPFC Control** | 0 | | | |
| | **dlPFC Control** | **vmPFC Control** | 0 | | | |
| | **Sham Stress** | **dlPFC Stress** | 0 | | | |

| | | | | | | |
|---|---|---|---|---|---|---|
| | **Sham Stress** | **vmPFC Stress** | 0 | | | |
| | **dlPFC Stress** | **vmPFC Stress** | 0 | | | |
| **Alpha Activity** | **Sham Control** | **Sham Stress** | 0 | | | |
| | **dlPFC Control** | **dlPFC Stress** | 0 | | | |
| | **vmPFC Control** | **vmPFC Stress** | 0 | | | |
| | **Sham Control** | **dlPFC Control** | 0 | | | |
| | **Sham Control** | **vmPFC Control** | 0 | | | |
| | **dlPFC Control** | **vmPFC Control** | **1** | **103.9** | **0.02** | **182 – 598** |
| | **Sham Stress** | **dlPFC Stress** | 0 | | | |
| | **Sham Stress** | **vmPFC Stress** | 0 | | | |
| | **dlPFC Stress** | **vmPFC Stress** | **1** | **119.41** | **0.02** | **154 – 628** |

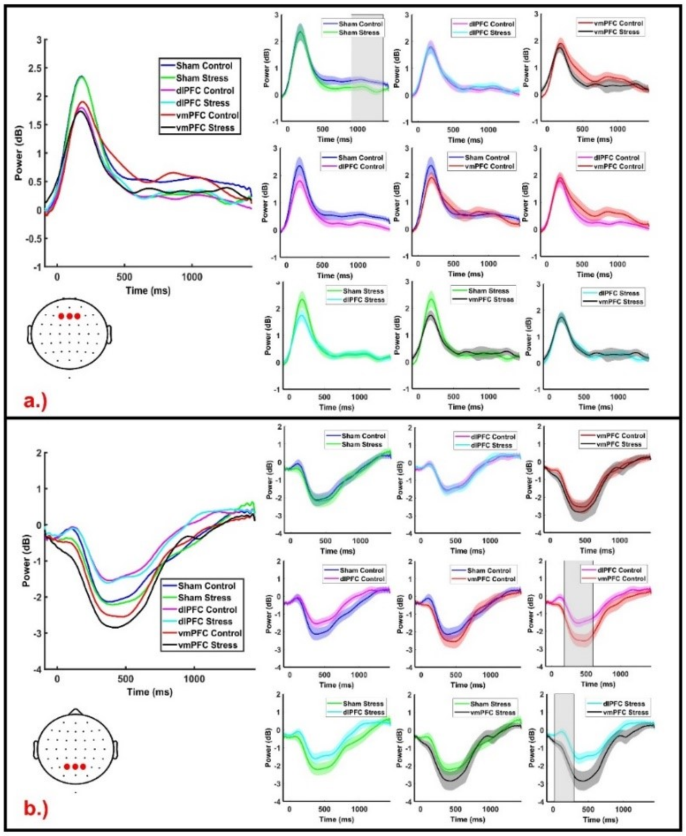


*Supplementary Figure 6 shows Time-frequency plots for task-related theta and alpha activity during the 3-Back task. a.) Theta ERD/ERS activity derived from frontal electrodes (Fz, F1, and F2) is shown with power indicated on the y-axis and time (in ms) on the x-axis in the left figure with averaged theta (4-8 Hz) activity over the time course for different conditions plotted together. The cartoon head below shows the positions of the target electrodes. The right side of the figures shows difference plots between conditions, as*

*indicated in the legend, with the shaded colored zones being the standard deviation for each condition. Cluster-based permutation testing was applied to identify significant clusters of time bins in the analysis. Significant clusters are highlighted by a grey box. b.) Alpha ERD/ERS activity from parietal electrodes (Pz, P1, and P2) are shown with power indicated by the y-axis and time (in ms) by the x-axis, plotted as described above with significant clusters highlighted in grey.*

**2.9 EEG Results (4-Back):** We analyzed the averaged P200 ERP component, P300 ERP component, task-based Theta ERS, and task-based Alpha ERD values (see supplementary Fig S6, Table S10).

**2.9.1 P200 ERP Component:** We found no significant main or interaction effects of emotional condition or stimulation group on the P200 ERP component.

**2.9.2 P300 ERP Component:** We found no significant main or interaction effect of emotional condition and stimulation group for the task-based P300 ERP component.

**2.9.3 Theta ERS:** We found no significant main or interaction effects of emotional condition and stimulation group for the task-based theta ERS activity.

**2.9.4 Alpha ERD:** We found no significant main or interaction effects of emotional condition and stimulation group for the task-based alpha ERD activity.

*Supplementary Table 11 Main and interaction effects of the ANOVAs conducted for EEG indexes for 4-Back load (P200 component, P300 component, Theta ERS, and Alpha ERD values). Significant p-values are marked in bold. d.f. = degrees of freedom, $\eta_p$2= partial eta squared.*

| | **Factors** | **d.f., Error** | **F Value** | **P - value** | $\eta_p^2$ |
|---|---|---|---|---|---|
| **P200** | **Emotional Condition** | 1,72 | .74 | .39 | .01 |
| **ERP (Cz)** | **Stimulation Group** | 2,72 | 1.86 | .16 | .049 |
| | **Emotional Condition x Stimulation Group** | 2,72 | .014 | .987 | .00 |
| **P300** | **Emotional Condition** | 1,72 | .24 | .624 | .003 |
| **ERP (Pz)** | **Stimulation Group** | 2,72 | .51 | .6 | .014 |
| | **Emotional Condition x Stimulation Group** | 2,72 | 1.94 | .15 | .051 |
| **Theta** | **Emotional Condition** | 1,72 | 1.8 | .185 | .024 |
| **ERS** | **Stimulation Group** | 2,72 | 2.67 | .076 | .069 |
| | **Emotional Condition x Stimulation Group** | 2,72 | .287 | .75 | .008 |

| | | | | | |
|---|---|---|---|---|---|
| **Alpha** | **Emotional Condition** | 1,72 | .055 | .816 | .001 |
| **ERD** | **Stimulation Group** | 2,72 | 2.01 | .141 | .053 |
| | **Emotional Condition x Stimulation Group** | 2,72 | .31 | .735 | .009 |

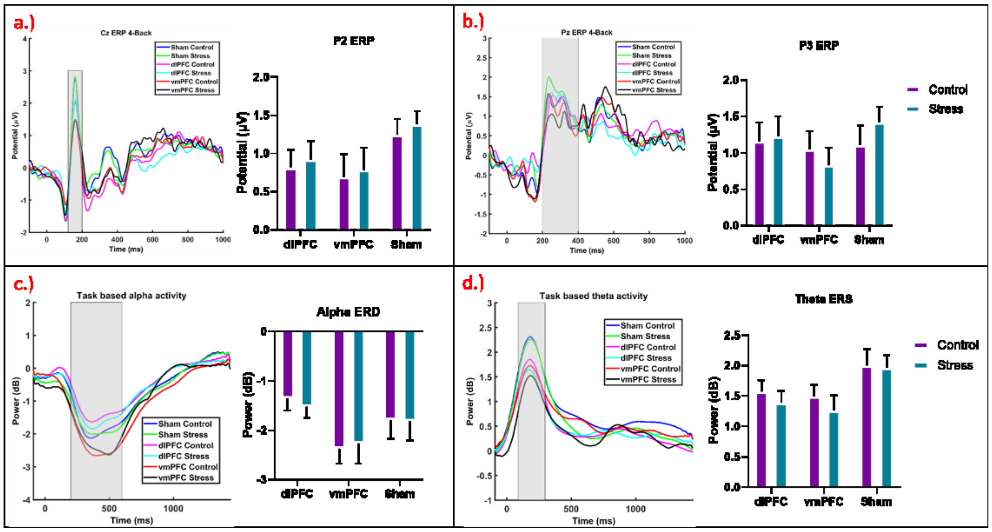


*Supplementary Figure 7 EEG results for the 4-Back load WM condition. a.) averaged ERP for all groups and conditions for EEG channel Cz. The x-axis represents time (in ms) and the y-axis shows the amplitude (µV). The P200 (120-200ms) component is highlighted by a gray box from which the averaged amplitude values were used for statistical analysis shown next to it in the bar plot. The bar plot shows the averaged P200 component values for all groups and conditions with x-axis delineating groups and the y-axis showing the potential amplitude (µV). b.) averaged ERP for all groups and conditions for the EEG channel Pz to show the P300 (200-400ms) component amplitude plotted as for P2. c.) averaged task-based alpha activity from parietal channels (Pz, P1, and P2) for all groups and conditions with the x-axis representing time (ms) and the y-axis power (in dB). The time range of interest for ERD (200-600 ms) is highlighted in grey from which averaged values are plotted in a bar graph with the x-axis delineating groups, and the y-axis power (dB). d.) averaged task-based theta activity from frontal channels (Fz, F1, and F2) for ERS (80-300ms) plotted as for alpha ERD. Error bars denote ± SEM.*

**2.10 TF Cluster Results (4-back):** We analyzed channel-averaged task-based theta and alpha activity over all 4-back epochs between conditions using Montecarlo-based permutation statistics with a cluster correction, as explained for the 2-Back load. Please refer to Table S11 for cluster information.

**2.10.1 Theta Activity:** No significant difference clusters were found between emotional conditions or stimulation groups in the 4-back load WM task for theta activity (see supplementary Figure 7a).

**2.10.2 Alpha Activity:** In the 4-back WM task condition, we only found significant differences between vmPFC and dlPFC groups in the control condition. We found a positive cluster from 154 ms to 618 ms where alpha ERD was larger in the vmPFC group as compared to the dlPFC group (see supplementary Figure 7b).

*Supplementary Table 12 Cluster information for the TF analysis of the 4-back load for task-based Alpha activity. The number of clusters, cluster t-values, p-values, and time range in each cluster are given for each analysis. The sign of the t value denotes the direction of effects, e.g. for condition 1 vs condition 2 for each analysis, a positive cluster shows a higher amplitude in condition 1, and a negative cluster shows a larger amplitude in condition 2. Only significant clusters are shown. Cluster t-values were calculated via cluster-based permutation testing using the Fieldtrip toolbox in EEGLAB.*

| | Condition 1 | Condition 2 | Number of Clusters | t-value | P-value | Time range |
|---|---|---|---|---|---|---|
| **Alpha Activity** | **Sham Control** | **Sham Stress** | 0 | | | |
| | **dlPFC Control** | **dlPFC Stress** | 0 | | | |
| | **vmPFC Control** | **vmPFC Stress** | 0 | | | |
| | **Sham Control** | **dlPFC Control** | 0 | | | |
| | **Sham Control** | **vmPFC Control** | 0 | | | |
| | **dlPFC Control** | **vmPFC Control** | **1** | **119.41** | **0.02** | **154 - 618** |
| | **Sham Stress** | **dlPFC Stress** | 0 | | | |
| | **Sham Stress** | **vmPFC Stress** | 0 | | | |
| | **dlPFC Stress** | **vmPFC Stress** | 0 | | | |

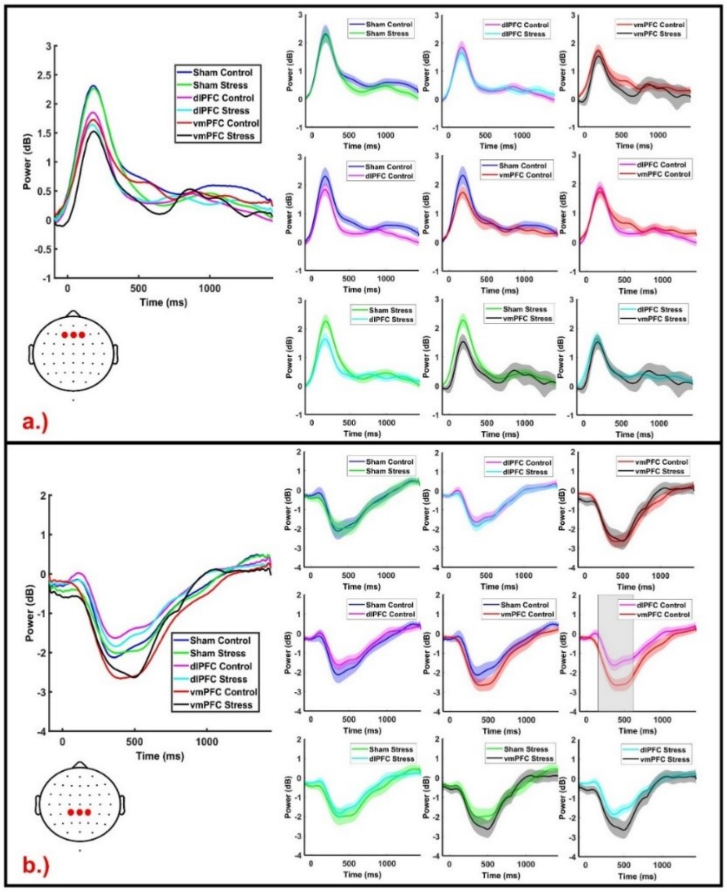


*Supplementary Figure 8 shows time-frequency plots for task-related theta and alpha activity during 4-Back task performance. a.) Theta ERD/ERS activity derived from frontal electrodes (Fz, F1, and F2) is shown with power on the y-axis and time (in ms) on the x-axis of the left figure with averaged theta (4-8 Hz) activity over the time course for different conditions plotted together. The cartoon head below shows the positions of the target electrodes. The right side of the figures shows difference plots between conditions, as indicated in the legend. The shaded colored zones indicate the standard deviation for each condition. Cluster-based permutation testing was applied to identify significant clusters of time bins in the analysis. b.) Alpha ERD/ERS activity from parietal electrodes (Pz, P1, and P2) are shown with power indicated by the y-axis and time (in ms) by the x-axis, plotted as described above with significant clusters highlighted in grey.*

**2.11 EEG Power –** We also analyzed EEG power changes caused by WM task performance and stimulation conditions by comparing power changes between the resting states RS2 and RS3 for theta and alpha power. The rmANOVAs performed for theta and alpha frequency bands for EO and EC states revealed a common pattern of effects, with a significant main effect of RS Block in all analyses revealing a power decrease for all groups, while no significant main or interaction effect of stimulation was found. Please refer to Table S12 for the results of the rmANOVAs (see Figure S8).

*Supplementary Table 13 Main and interaction effects of the ANOVAs conducted for power in the theta and alpha ranges in resting state (RS) blocks (theta eyes open, theta eyes closed, alpha eyes open, and alpha eyes closed) are shown. Significant p-values are marked in bold, d.f. = degrees of freedom, $\eta_p 2$= partial eta squared.*

| | Factors | d.f., Error | F Value | P - value | $\eta_p^2$ |
|---|---|---|---|---|---|
| **Theta Eyes Open** | **Emotional Condition** | 1, 75 | 2.144 | 0.124 | 0.031 |
| | **RS Block** | 1, 75 | 5.55 | **0.021*** | 0.069 |
| | **Stimulation Group** | 2, 75 | 0.107 | 0.898 | 0.003 |
| | **Emotional Condition x RS Block** | 1, 75 | 0.383 | 0.538 | 0.005 |
| | **Emotional condition x Stimulation Group** | 2, 75 | 0.589 | 0.558 | 0.015 |
| | **Stimulation group x RS Block** | 2, 75 | 0.27 | 0.764 | 0.007 |
| | **Emotional Condition x Time x Stimulation** | 2, 75 | 1.107 | 0.336 | 0.029 |
| **Theta Eyes Closed** | **Emotional Condition** | 1, 75 | 3.81 | 0.055 | 0.048 |
| | **RS Block** | 1, 75 | 13.19 | **<0.001*** | 0.15 |
| | **Stimulation Group** | 2, 75 | 1.21 | 0.304 | 0.031 |
| | **Emotional Condition x RS Block** | 1, 75 | 0.014 | 0.906 | 0.000 |
| | **Emotional condition x Stimulation Group** | 2, 75 | 0.162 | 0.851 | 0.004 |
| | **Stimulation group x RS Block** | 2, 75 | 0.903 | 0.410 | 0.024 |
| | **Emotional Condition x Time x Stimulation** | 2, 75 | 0.029 | 0.971 | 0.001 |
| **Alpha Eyes Open** | **Emotional Condition** | 1, 75 | 0.02 | 0.888 | 0.000 |
| | **RS Block** | 1, 75 | 11.255 | **0.001*** | 0.13 |
| | **Stimulation Group** | 2, 75 | 0.328 | 0.721 | 0.009 |
| | **Emotional Condition x RS Block** | 1, 75 | 0.069 | 0.794 | 0.001 |
| | **Emotional condition x Stimulation Group** | 2, 75 | 0.993 | 0.375 | 0.026 |
| | **Stimulation group x RS Block** | 2, 75 | 0.526 | 0.593 | 0.014 |
| | **Emotional Condition x Time x Stimulation** | 2, 75 | 2.23 | 0.115 | 0.056 |
| **Alpha Eyes Closed** | **Emotional Condition** | 1, 75 | 0.498 | 0.483 | 0.007 |
| | **RS Block** | 1, 75 | 11.99 | **<0.001*** | 0.138 |
| | **Stimulation Group** | 2, 75 | 2.032 | 0.138 | 0.051 |
| | **Emotional Condition x RS Block** | 1, 75 | 0.944 | 0.334 | 0.012 |
| | **Emotional condition x Stimulation Group** | 2, 75 | 0.028 | 0.972 | 0.001 |
| | **Stimulation group x RS Block** | 2, 75 | 0.412 | 0.664 | 0.011 |
| | **Emotional Condition x Time x Stimulation** | 2, 75 | 0.057 | 0.945 | 0.002 |

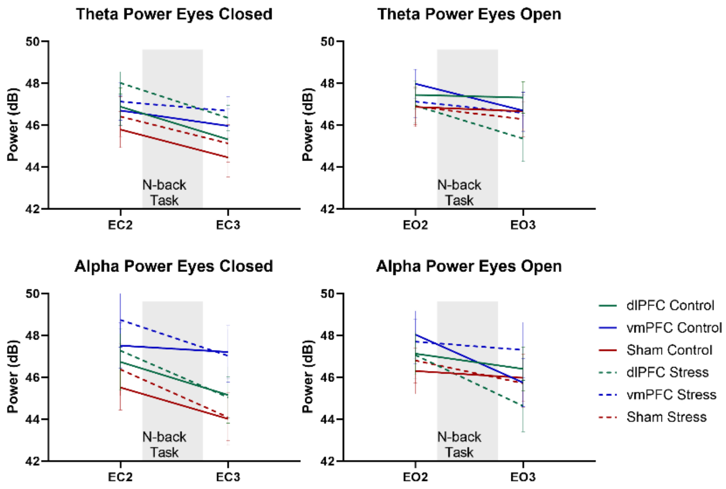


*Supplementary Figure 9 EEG power changes in eyes open and eyes closed conditions for the theta (4-8 Hz) frequency range obtained from frontal electrodes (Fz, F1, F2) and the alpha (8-13 Hz) frequency range obtained from parietal electrodes (Pz, P1, P2). The electrodes of interest were identical to the task-related TF analysis. The y-axis denotes the power in decibels and the x-axis shows the resting state blocks before and after task performance. No significant differences between stimulation groups and conditions were identified. Error bars show the Standard Error of Mean (SEM).*

**2.12 Correlations –** We also performed multiple correlation analyses to understand the association of EEG parameters, and other physiological indices on Hits, and D-Prime scores for each stimulation group and emotional condition separately.

***2.12.1 2-Back:*** In the 2-back load condition, we identified a significant positive correlation of cortisol levels after task performance with D-prime scores for the vmPFC stress condition (see Figure S9). No other significant correlations were found.

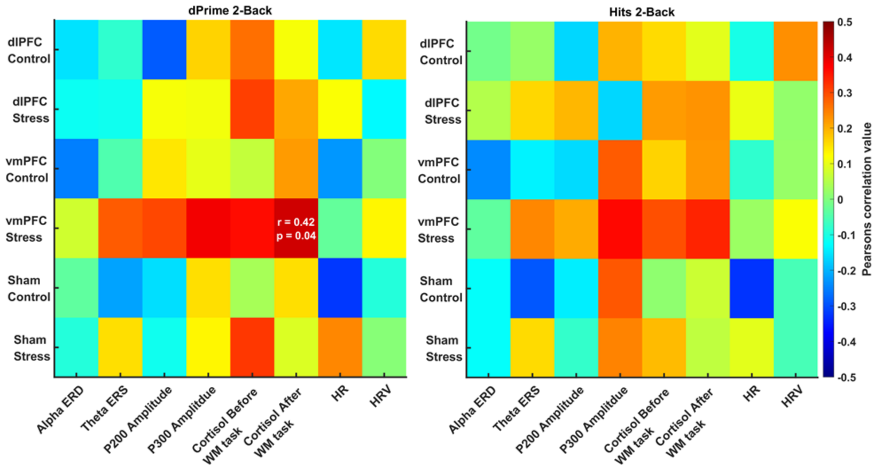


*Supplementary Figure 10 Heat map for Pearson's correlation values in the 2-back load WM task for the EEG (alpha ERD, theta ERS, P200 amplitude, P300 amplitude) and physiological indexes (cortisol levels before and after task performance, HR, and HRV during task performance) with the performance parameters Hits and D-prime for all stimulation and emotional conditions. The colors of the plot are showing r values with the color bar in the range from 0.5 to -0.5 with red showing positive and blue showing negative correlations. For significant correlations, r-values, and p-values of respective correlations are reported.*

***2.12.2 4-Back:*** In the 4-back load condition, we found a significant negative correlation of the P200 with both hits and D-prime scores in the sham control conditions. Theta ERS also negatively correlated with Hits in the sham Control condition, and HR negatively correlated with D-prime in the sham control conditions. Theta ERS was positively correlated with D-prime in the vmPFC control condition and with hits in the dlPFC stress condition. Similar to the 2- and 3-back load conditions, cortisol levels after task performance were correlated positively with D-prime scores in the vmPFC stress condition. The P300 amplitude correlated positively with the sham stress condition (see figure S10).

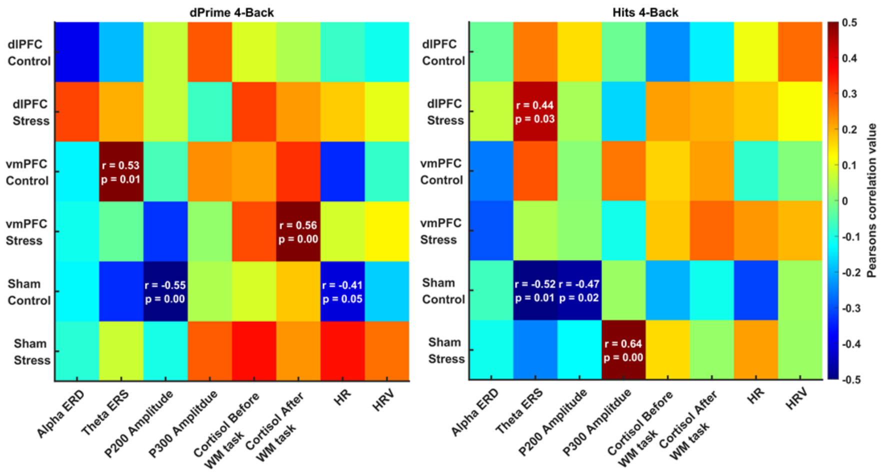


*Supplementary Figure 11 Heat map for Pearson correlation values in the 4-back load WM task for the EEG (alpha ERD, theta ERS, P200 amplitude, P300 amplitude) and physiological indexes (cortisol levels before and after task performance, HR, and HRV during task performance) with the performance parameters Hits and D-prime for all stimulation and emotional conditions. The colors of the plot are showing r values with the color bar in the range from 0.5 to -0.5 with red showing positive and blue showing negative correlations. For significant correlations, r-values, and p-values are reported.*
